# Population structure, biogeography and transmissibility of *Mycobacterium tuberculosis*

**DOI:** 10.1101/2020.09.29.293274

**Authors:** Luca Freschi, Roger Vargas, Ashek Hussain, S M Mostofa Kamal, Alena Skrahina, Sabira Tahseen, Nazir Ismail, Anna Barbova, Stefan Niemann, Daniela Maria Cirillo, Anna S Dean, Matteo Zignol, Maha Reda Farhat

**Affiliations:** Department of Biomedical Informatics, Harvard Medical School, Boston, MA, USA; Department of Systems Biology, Harvard Medical School, Boston, MA, USA; Pulmonary and Critical Care Medicine, Massachusetts General Hospital, Boston, MA, USA; Directorate General of Health Services, Ministry of Health and Family Welfare, Dhaka, Bangladesh; Department of Pathology and Microbiology, National Institute of Diseases of the Chest and Hospital, Dhaka, Bangladesh; Republican Scientific and Practical Centre for Pulmonology and Tuberculosis, Minsk, Belarus; National Reference Laboratory, National Tuberculosis Control Programme, Islamabad, Pakistan; National Institute for Communicable Diseases, Sandringham, South Africa; Department of Medical Microbiology, University of Pretoria, Pretoria, South Africa; Central Reference Laboratory on Tuberculosis Microbiological Diagnostics, Ministry of Health, Kiev, Ukraine; Molecular and Experimental Mycobacteriology, Borstel Research Centre, Borstel, Germany; Emerging Bacterial Pathogens Unit, IRCCS San Raffaele Scientific Institute, Milan, Italy; Global Tuberculosis Programme, World Health Organization, Geneva, Switzerland

## Abstract

*Mycobacterium tuberculosis* is a clonal pathogen proposed to have co-evolved with its human host for millennia, yet our understanding of its genomic diversity and biogeography remains incomplete. Here we use a combination of phylogenetics and dimensionality reduction to reevaluate the population structure of *M. tuberculosis*, providing the first in-depth analysis of the ancient East African Indian Lineage 1 and the modern Central Asian Lineage 3 and expanding our understanding of Lineages 2 and 4. We assess sub-lineages using genomic sequences from 4,939 pan-susceptible strains and find 30 new genetically distinct clades that we validate in a dataset of 4,645 independent isolates. We characterize sub-lineage geographic distributions and demonstrate a consistent geographically restricted and unrestricted pattern for 20 groups, including three groups of Lineage 1. We assess the transmissibility of the four major lineages by examining the distribution of terminal branch lengths across the *M. tuberculosis* phylogeny and identify evidence supporting higher transmissibility in Lineages 2 and 4 than 3 and 1 on a global scale. We define a robust expanded barcode of 95 single nucleotide substitutions (SNS) that allows for the rapid identification of 69 *Mtb* sub-lineages and 26 additional internal groups. Our results paint a higher resolution picture of the *Mtb* phylogeny and biogeography.

## Introduction

Tuberculosis (TB) is among the ten top causes of death worldwide. In 2018 more than ten million people fell ill and 1.5 million died from TB (https://www.who.int/tb/global-report-2019). During the last two decades significant efforts have been made to understand strain level genetic diversity in the TB bacillus *Mycobacterium tuberculosis* (*Mtb*) and its geographic distribution. A robust classification of *Mtb* strains into evolutionarily meaningful sub-lineages is important both for taxonomic purposes and because sub-lineages can differ in virulence or antibiotic resistance ^1^. *Mtb* was first classified into three principal genetic groups in 1997, based on two neutral single nucleotide substitutions (SNSs) in the antibiotic resistance genes *katG* (codon 463) and *gyrA* (codon 95) ^2^. Since then several studies have attempted higher resolution classification, using large genomic deletions ^3^, spoligotyping ^4^ and SNSs ^5–11^. There are currently eight recognized *Mtb* lineages (L1-8), although only two lineages (L2, L4) have been well represented in taxonomic and phylogeographic evaluations ^8–12^. Within the major lineages, 53 sub-lineages have been described and are definable with an SNS ‘barcode’ ^8^: 7 for L1, 6 for L2, 4 for L3 and 36 for L4. These 53 sub-lineages best characterize diversity within L2 and L4. L1 and L3 diversity is less understood as these lineages are most prevalent in countries where pathogen sequencing has been less widely applied, but this is rapidly changing due to the increasing sequencing capacity in high-burden TB settings and supported by international research collaborations ^13^.

Host-pathogen co-evolution and more recent host-related selective pressures have been postulated to drive genetic diversity in *Mtb* ^14,15^. *Of high relevance to public health, this genetic diversity is thought to underlie the observed differences in Mtb* sub-lineage transmissibility ^16,17^ or host specificity ^10^. For example, of the *Mtb* sub-lineages prevalent in Ho Chi Minh City, Vietnam, one modern L2 sub-lineage (2.2.1) was found to be more transmissible relative to other L2 sub-lineages, L4 or L1 ^16^. The phylogeography of L5 and L6 (also known as *M. africanum)*, L7, L8 and even some L4 sub-lineages (*e*.*g*. 4.6.1/Uganda, 4.6.2/Cameroon, 4.1.3/Ghana and 4.5) has demonstrated that these groups are more geographically restricted than L2 and other L4 sub-lineages. These geographically restricted groups are associated with lower variation in *Mtb* T-cell epitopes and support the notion that *Mtb* sub-lineages have a spectrum of pathogenic strategies and may be niche specialists infecting preferentially humans of a specific population or ancestry ^10^. However, more evidence is needed to support this hypothesis given the relatively few genetic differences that define *Mtb* sub-lineages or human ancestry.

Here, we sought to expand our understanding of the *Mtb* population structure by leveraging 9,584 genomes sampled from 49 countries, including 738 L1 and 1,104 L3 isolates. We identify and validate 22 novel sub-lineages and 8 additional internal groups (*i*.*e*. genetically divergent groups found in sub-lineages that cannot be further partitioned in a hierarchical fashion according to our criteria), including 6 in L1 and 4 in L3, and expand the SNS typing barcode to 95 sites. We find that L3 and L2 have similar phylogenetic structure, yet L2 along with L4 manifest a phylogenetic signal of increased transmissibility compared with L1 or L3. These findings, along with our observations of novel geographically restricted sub-lineages, expand the evidence supporting the *Mtb* human co-evolution hypothesis.

## Results

### Detailed population structure of L1-4 and a hierarchical sub-lineage naming system

We assembled a high-quality dataset of whole genomes, antibiotic resistance phenotypes and geographic sites of isolation for 9,584 clinical *Mtb* samples (Methods and Suppl. File 1). Of the total, 4,939 (52%) were pan-susceptible, *i*.*e*. susceptible to at least isoniazid and rifampicin (and all other antibiotics when additional phenotypic data were available), and 4,645 (48%) were resistant to one or more antibiotics (Suppl. Fig. 1a). Using the 62 SNS lineage barcode ^8^, 738 isolates were classified as L1 (8%), 2,193 as L2 (22%), 1,104 as L3 (12%) and 5,549 as L4 (58%, Suppl. Fig. 1b). Among the 4,939 pan-susceptible isolates, we identified high-quality genome-wide SNSs (83,735 for L1, 56,736 for L2, 76,817 for L3 and 185,622 for L4) that we used in building maximum-likelihood phylogenies for each major lineage (L1-4, Methods). We computed an index of genetic divergence (F_ST_) between groups defined by each bifurcation in each phylogeny. Sub-lineages were defined as monophyletic groups that had high F_ST_ (> 0.33) and were also clearly separated from other groups in principal component analysis (PCA, Methods). We also defined internal groups to sub-lineages (Methods): an internal group is a monophyletic group genetically divergent (by F_ST_ and PCA) from its neighboring groups, but has one or more ancestral branches that show a low degree of divergence or low support (bootstrap values). Internal groups do not represent true sub-lineages in a hierarchical fashion, but defining them allows us to further characterize the *Mtb* population structure. We provide code to automate all the steps described above. Our approach is scalable and can be used on other organisms (Methods).

**Fig. 1.**
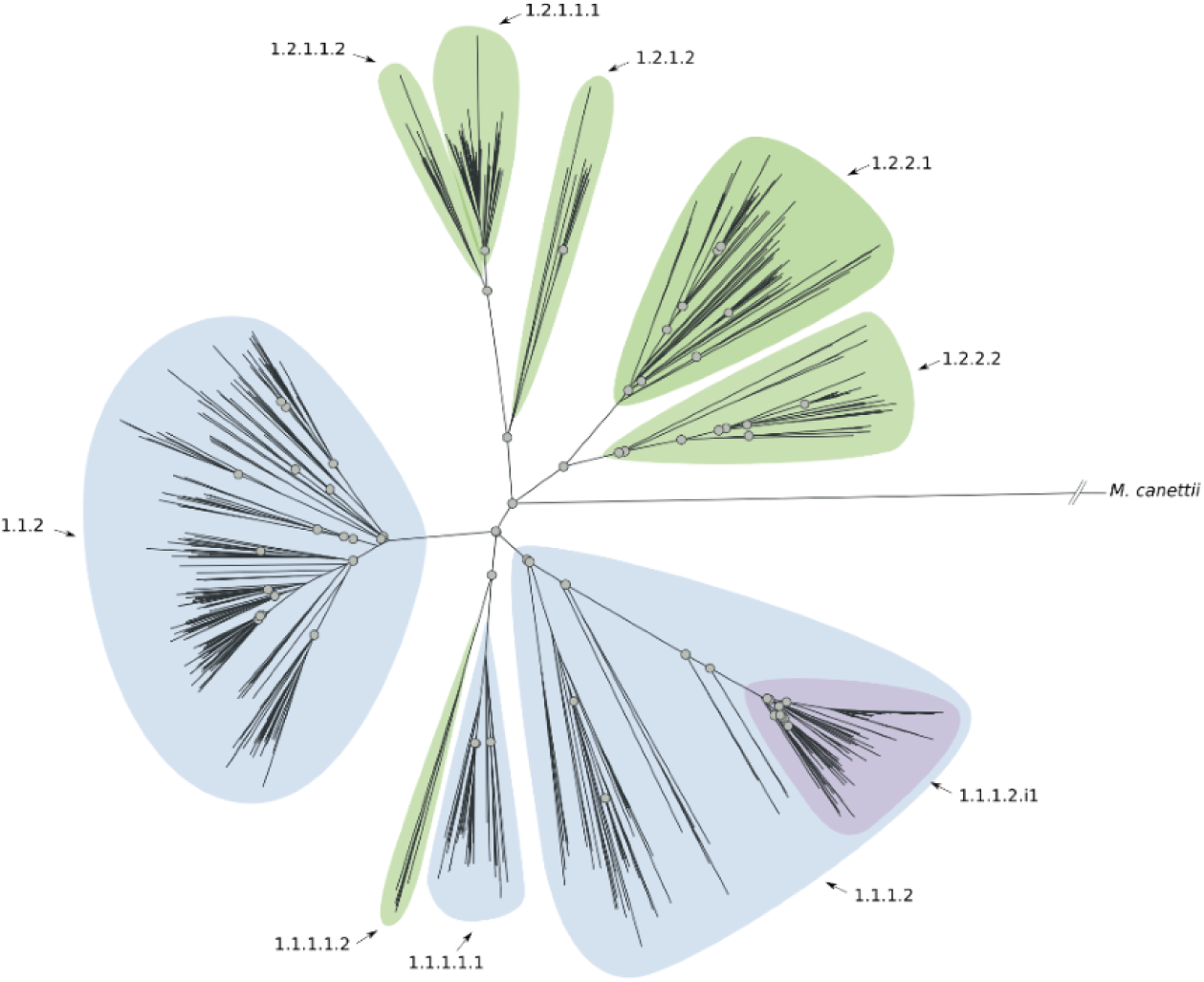
Phylogenetic tree reconstruction of L1 (binary tree). Grey circles define splits where the F_ST_ calculated using the descendants of the two children nodes is greater than 0.33. The sub-lineages are defined by colored areas (blue: sub-lineages already described in the literature; green: sub-lineages described here; purple: internal sub-lineages).

To better classify *Mtb* isolates in the context of the global *Mtb* population structure, we developed a hierarchical sub-lineage naming scheme (Suppl. File 2) where each subdivision in the classification corresponds to a split in the phylogenetic tree of each major *Mtb* lineage. Starting with the global *Mtb* lineage numbers (*e*.*g*., L1), we recursively introduced a subdivision (*e*.*g*., from 1.2 to 1.2.1 and 1.2.2) at each bifurcation of the phylogenetic tree whenever both subclades sufficiently diverged. Formally, we defined these splits using bootstrap criteria, and independent validations by F_ST_ and PCA (Methods). Internal groups were denoted with the letter “i” (*e*.*g*. 4.1.i1). This proposed system overcomes two major shortcomings of the existing schemas: same-level sub-lineages are never overlapping (unlike the system of Stucki *et al*. ^10^ sub-lineage 4.10 includes sub-lineages 4.7–4.9), and the names reflect both phylogenetic relationships and genetic similarity (unlike semantic naming such as the “Asia ancestral” lineage in the system of Shitikov *et al*. ^9^).

Using the sub-lineage definition rules and the sub-lineage naming scheme described above, we characterized six previously undescribed sub-lineages of L1 (Fig. 1 and Suppl. Fig. 2); five of which expand the current description of 1.2. We also detected an internal group of 91 isolates (1.1.1.2.i1) characterized by a long defining branch in the phylogeny (corresponding to 82 SNSs), a high F_ST_ (0.48), and geographically restricted to Malawi (85/91, 93% isolates). We found four previously undescribed sub-lineages of L3 (Fig. 2, Suppl. Fig. 3), revising L3 into four main groups, whereas previously only two partitions of one sub-lineage were characterized (3.1). We found that the latter two partitions are in fact internal groups of the largest sub-lineage (3.1.1) in our revised classification.

**Fig. 2.**
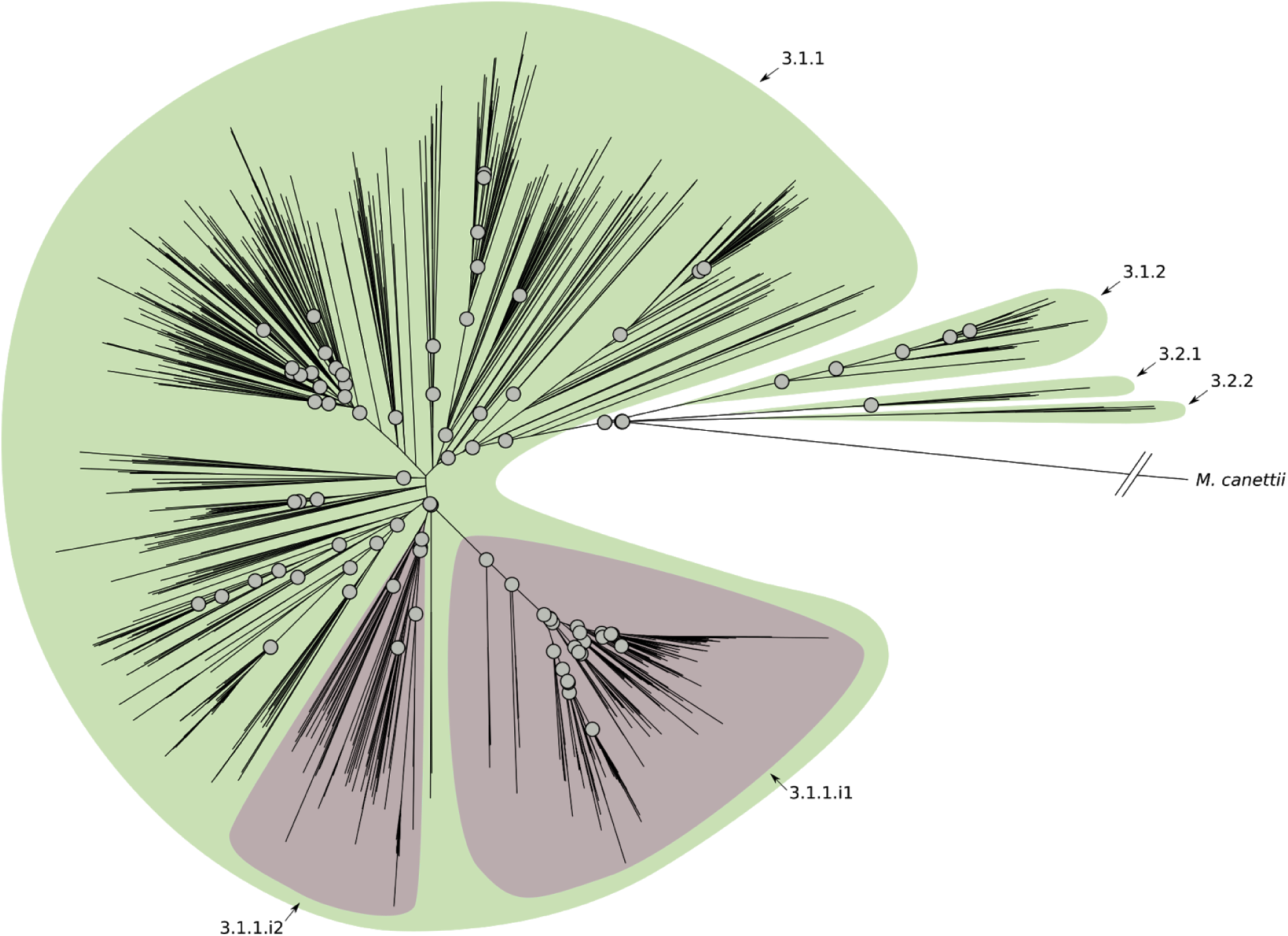
Phylogenetic tree reconstruction of L3 (binary tree). Grey circles define splits where the F_ST_ calculated using the descendants of the two children nodes is greater than 0.33. The sub-lineages are defined by colored areas (green: sub-lineages described here; purple: internal sub-lineages).

**Fig. 3.**
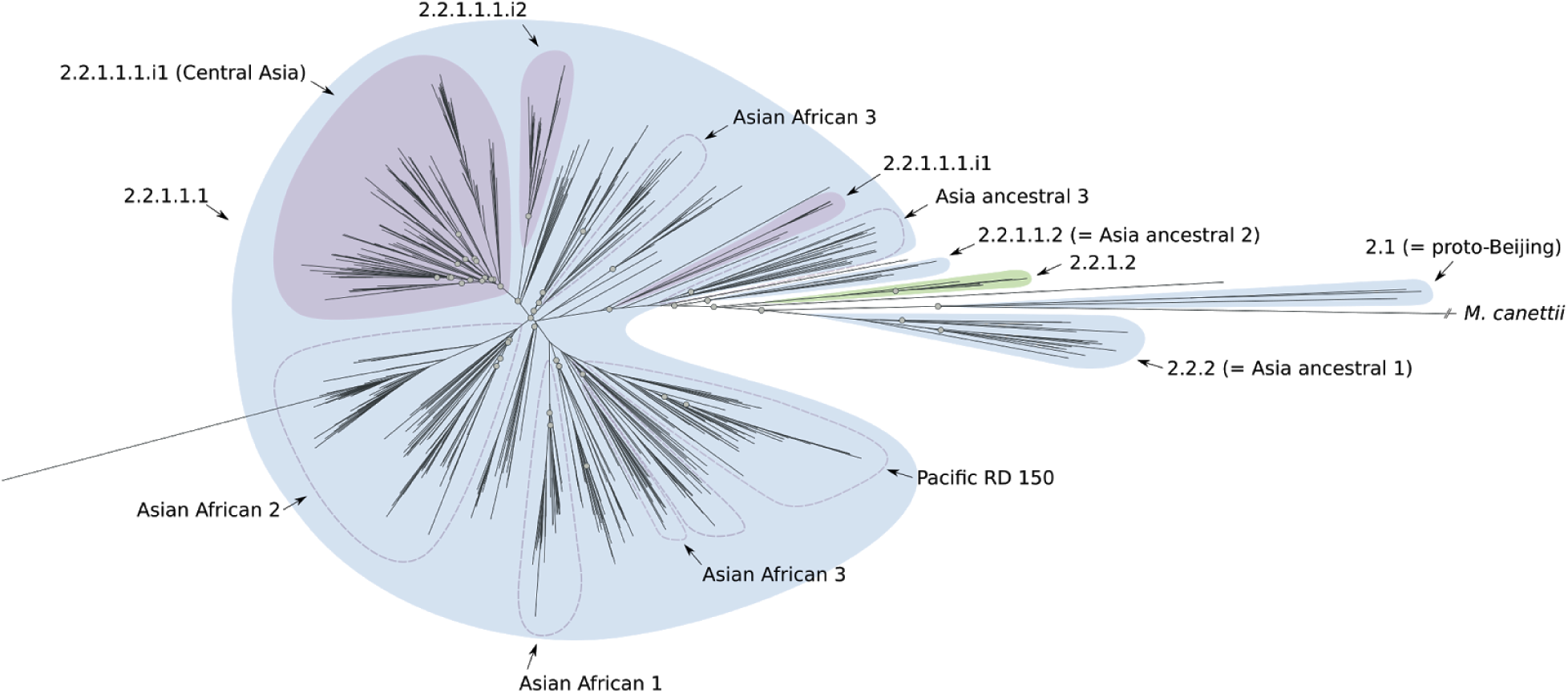
Phylogenetic tree reconstruction of L2 (binary tree). Grey circles define splits where the F_ST_ calculated using the descendants of the two children nodes is greater than 0.33. The sub-lineages are defined by colored areas (blue: sub-lineages already described in the literature; green: sub-lineages described here; purple: internal sub-lineages).

L2 is divided into two groups: proto-Beijing and Beijing with the latter in turn partitioned into two groups: ancient- and modern-Beijing ^9^. Each one of these groups is characterized by further subdivisions (three for the ancient-Beijing group and seven for the modern-Beijing group; see Suppl. Fig 4). We found a new sub-lineage (2.2.1.2, Fig. 3, Suppl. Fig. 4) within the previously characterized ancient-Beijing group. However, genetic diversity within the modern-Beijing group (2.2.1.1.1) was lower than in the other L2 sub-lineages and the tree topology and F_ST_ calculations did not support further hierarchical subdivisions. Although we did find three internal groups of modern-Beijing: two undescribed and one that corresponds to the Central Asia group ^9^. For L4, our results support a complex population structure with 21 sub-lineages and 15 internal groups. In particular, we found 11 previously undescribed sub-lineages and 5 internal groups that expand our understanding of previously characterized sub-lineages (*e*.*g*., 4.2.2; 4.2 in the Coll *et al*. classification) or that were not characterized since these isolates were simply classified as L4 (*e*.*g*., 4.2.1.1.1.1.2, Fig. 4, Suppl. Fig. 5) using the other barcodes.

**Fig. 4.**
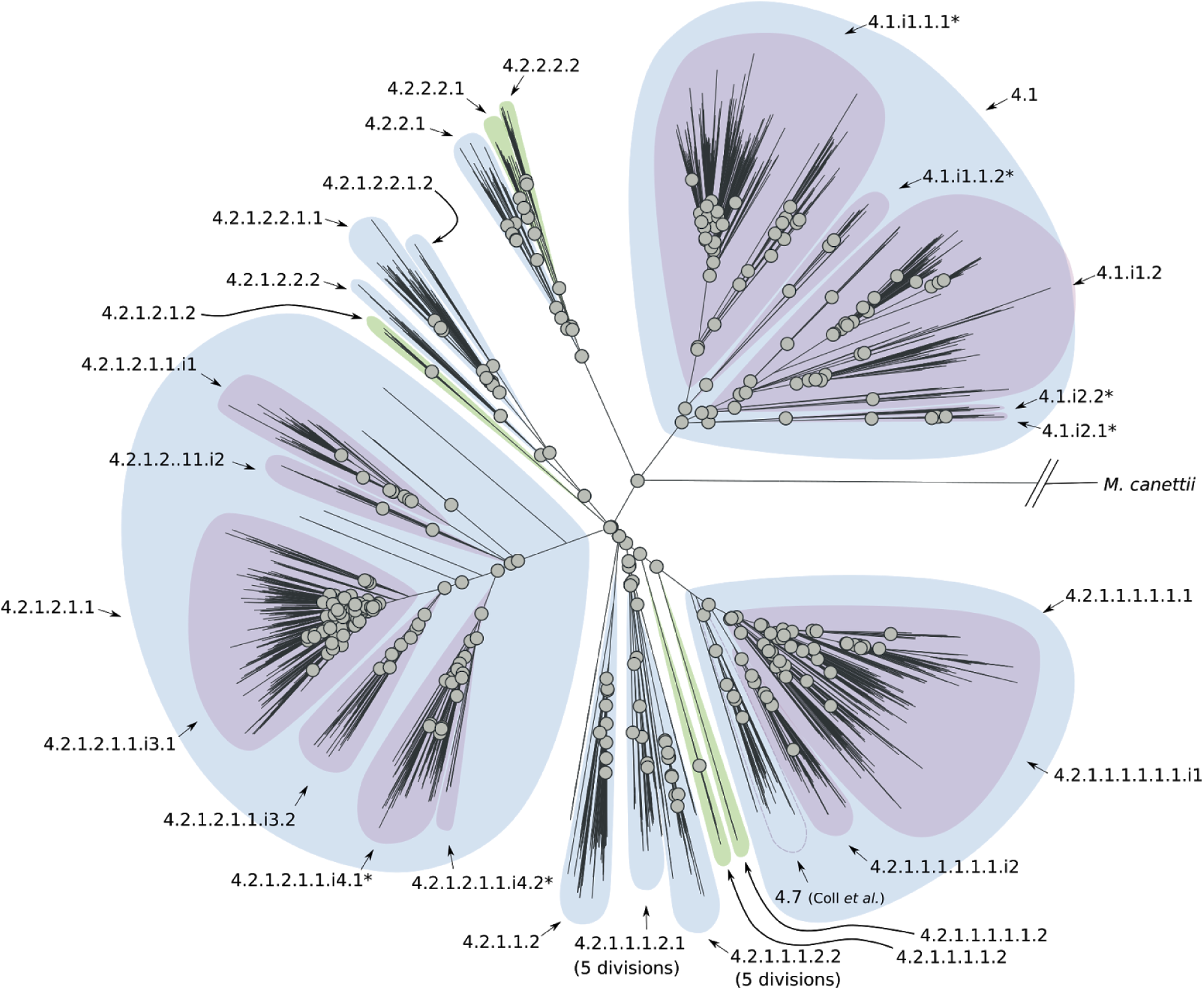
Phylogenetic tree reconstruction of L4 (binary tree). Grey circles define splits where the F_ST_ calculated using the descendants of the two children nodes is greater than 0.33. The sub-lineages are defined by colored areas (blue: sub-lineages already described in the literature; green: sub-lineages described here; purple: internal sub-lineages).

**Fig. 5.**
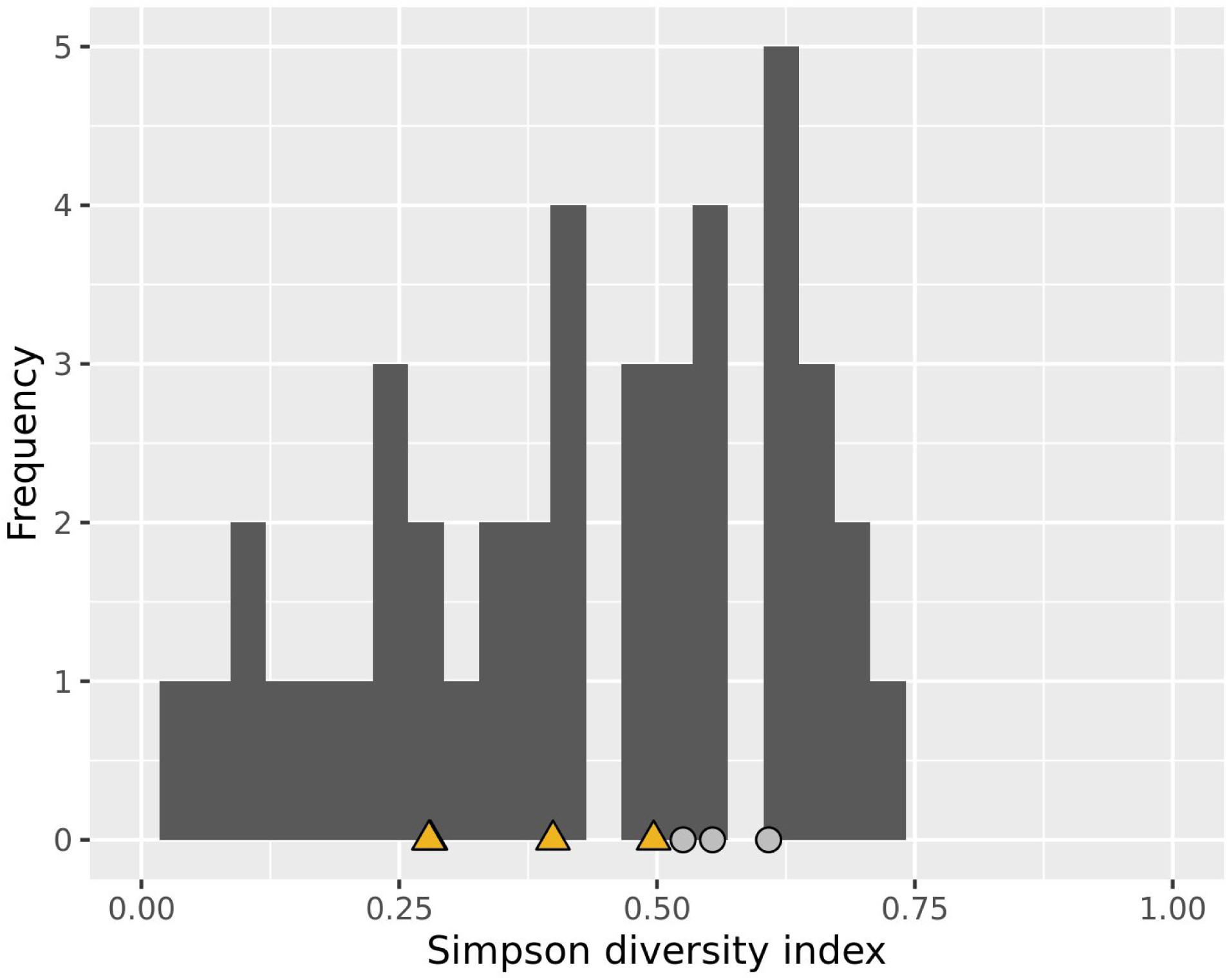
Histogram of the Simpson diversity index calculated for all sub-lineages of L1-4. A dataset of 17,432 isolates from 74 countries was used to perform this analysis. Yellow triangles designate the Simpson diversity index values of sub-lineages designated as geographically restricted by Stucki *et al*. Light grey circles designate the Simpson diversity index values of sub-lineages designated as geographically unrestricted by Stucki *et al*.

### A new barcode to define L1-4 *Mtb* sub-lineages and a software package to type *Mtb* strains from WGS data

We defined a SNS barcode for distinguishing the obtained sub-lineages (Suppl. File 3). We characterized new synonymous SNSs found in 100% of isolates from a given sub-lineage, but not in other isolates from the same major lineage, compiling 95 SNSs into an expanded barcode (Suppl. File 3). We validated the barcode by using it to call sub-lineages in the hold-out set of 4,645 resistant isolates and comparing the resulting sub-lineage designations with maximum likelihood phylogenies inferred from the full SNS data (Suppl. Fig. 6-9). A sub-lineage was validated if it was found in the hold-out data and formed a monophyletic group in the phylogeny. Considering the “recent” sub-lineages, *i*.*e*. the most detailed level of classification in our system, we were able to validate eight out of nine L1 sub-lineages including five out of six of the new sub-lineages described here, with the exception of 1.1.1.1.2. We validated all four new L3 sub-lineages, all five L2 sub-lineages including the one previously undescribed, and 16 of the 21 L4 sub-lineages including two described here. The sub-lineages we could not confirm were not represented by any isolate in the validation phylogenies. We did not observe any paraphyletic sub-lineages in the revised classification system.

**Fig. 6.**
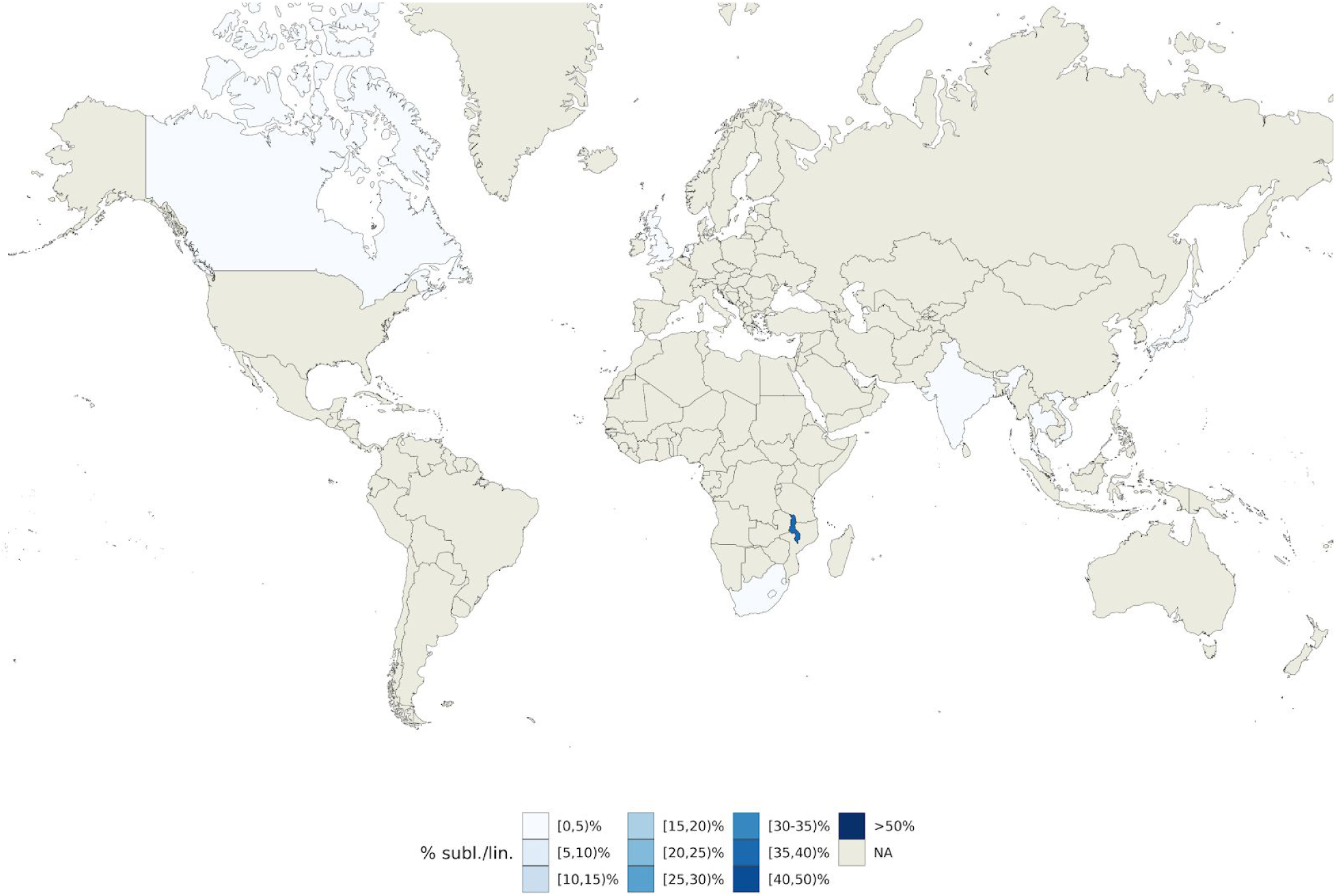
World map showing the geographic distribution of internal sub-lineage 1.1.1.2.i1. Colors represent the percentage of 1.1.1.2.i1 strains isolated in a given country with respect to all L1 strains isolated in such country.

We developed fast-lineage-caller, a software tool that classifies *Mtb* genomes using the SNS barcode proposed above. For a given genome, it returns the corresponding sub-lineage as output using our hierarchical naming system in addition to four other existing numerical/semantic naming systems, when applicable (Methods). The tool also informs the user on how many SNSs support a given lineage call and allows for filtering of low quality variants. The tool is generalizable and can manage additional barcodes defined by the user including for species other than *Mtb*.

### Geographic distribution of the *Mtb* sub-lineages

Next, we examined whether certain sub-lineages were geographically restricted, which would support the *Mtb*-human co-evolution hypothesis, or whether they constituted prevalent circulating sub-lineages in several different countries (*i*.*e*. geographically unrestricted) ^10^. We used our SNS barcode to determine the sub-lineages of 17,432 isolates (Methods) sampled from 74 countries (Suppl. Fig. 10, Suppl. File 4). We computed the Simpson diversity index (Sdi) as a measure of geographic diversity that controls for variable sub-lineage frequency (Methods) for each well-represented sub-lineage or internal group (n > 20). We hypothesized that geographically unrestricted lineages would have a higher Sdi. We found Sdi to correlate highly (ρ = 0.68; p-value = 5.7 × 10^−7^) with the number of continents from which a given sub-lineage was isolated (Suppl. Fig. 11). The Sdi ranged between a minimum of 0.05 and a maximum of 0.72, with a median value of 0.46 (Fig. 5). The known geographically restricted sub-lineages ^10^ had an Sdi between 0.28 and 0.5 (Fig. 5, Suppl. Table 1), while the known geographically unrestricted sub-lineages ^10,11^ had an Sdi between 0.55 and 0.61 (Fig. 5, Suppl. table 2). We found 11 sub-lineages/internal groups with Sdi <0.28 (Suppl. Table 3), and 11 sub-lineages/internal groups with Sdi >0.61 (Suppl. Table 4), *i*.*e*. more extreme than previously reported geographically restricted or unrestricted sub-lineages, respectively.

While the currently known geographically restricted sub-lineages are all in L4, we found evidence of geographic restriction for two sub-lineages/internal groups of L1. The first, the L1 internal group 1.1.1.2.i1, showed a very low Sdi (0.06) and was only found at high frequency among the circulating L1 isolates in Malawi (Fig. 6). This finding is also in agreement with the L1 phylogeny (Fig. 1) that shows a relatively long (82 SNS) branch defining this group. The second geographically restricted L1 sub-lineage is 1.1.1.1.1 (Sdi = 0.12) that was only found at high frequency among circulating L1 isolates in South-East Asia (Vietnam and Thailand, Fig. 7). To exclude the possibility that these two groups appeared geographically restricted as a result of oversampling transmission outbreaks, we calculated the distribution of the pairwise SNS distance for each of these two sub-lineages. We measured a median SNS distance of 204 and 401, respectively, refuting this kind of sampling error for these groups (typical pairwise SNS distance in outbreaks <15-20 SNS ^18^) (Suppl. Fig. 12).

**Fig. 7.**
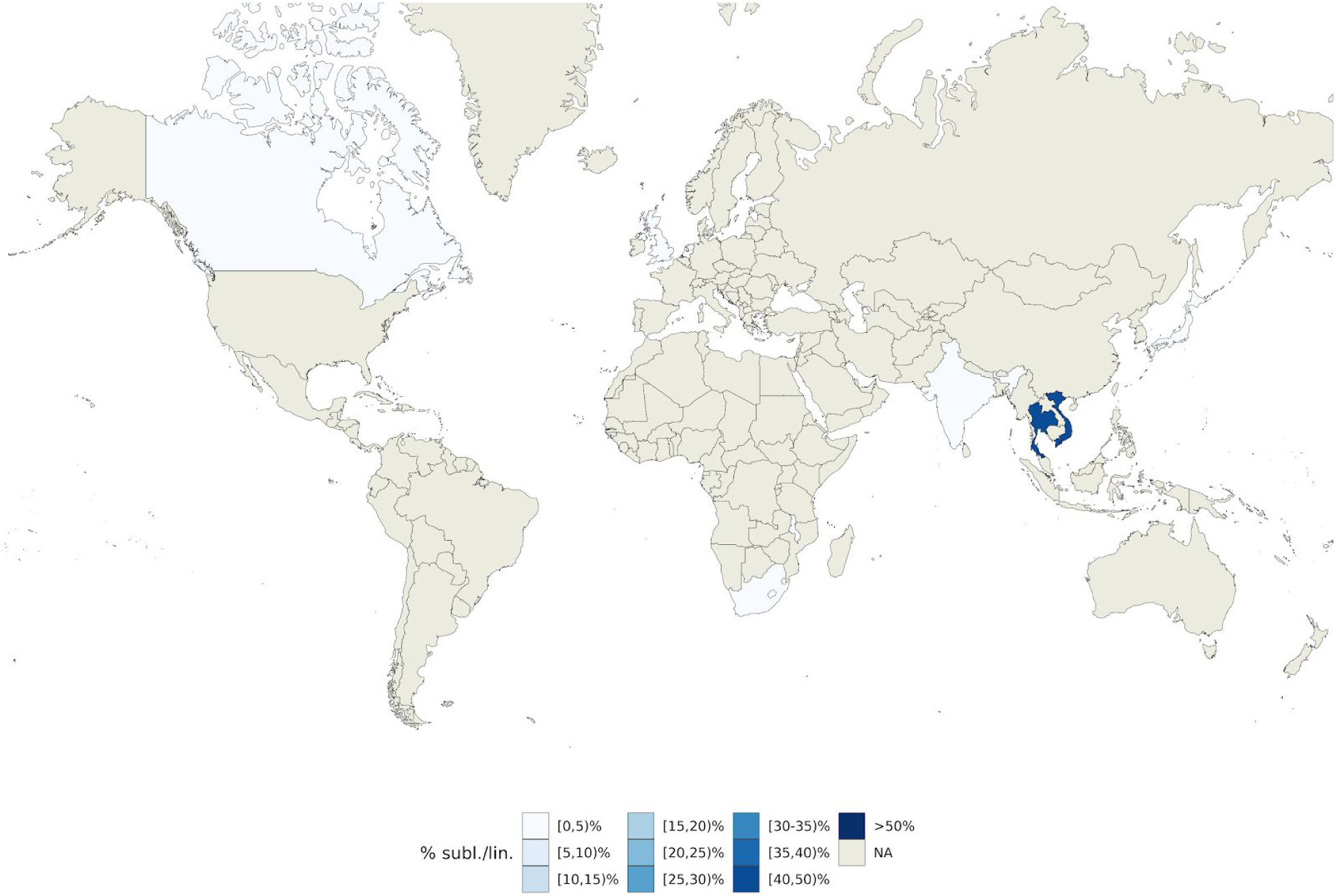
World map showing the geographic distribution of internal sub-lineage 1.1.1.1.1. Colors represent the percentage of 1.1.1.1.1 strains isolated in a given country with respect to all L1 strains isolated in such country.

The geographically unrestricted sub-lineages spanned several recognized generalist sub-lineages or internal groups (*e*.*g*. 4.1.i1.1.1.1, 4.2.1.2.1.1, 4.2.1.1.1.1.1.1 and 2.2.1.1.1, which correspond to 4.1.2/Haarlem, 4.3/LAM, 4.10/PGG3, modern-Beijing in the Stucki *et al*. ^*10*^ or the Shitikov *et al*. ^9^ classification). In addition, we found a candidate geographically unrestricted sub-lineage of L1 (1.1.2). L1.1.2 spanned 253 isolates from 7 countries and 4 continents and its Sdi was 0.61 (Fig. 8). Overall we found a low, but significant correlation between Sdi and the number of isolates sampled from each sub-lineage (ρ = 0.34; p-value = 0.03, Suppl. Fig. 13) indicating that the most prevalent sub-lineages were more likely to be geographically unrestricted.

**Fig. 8.**
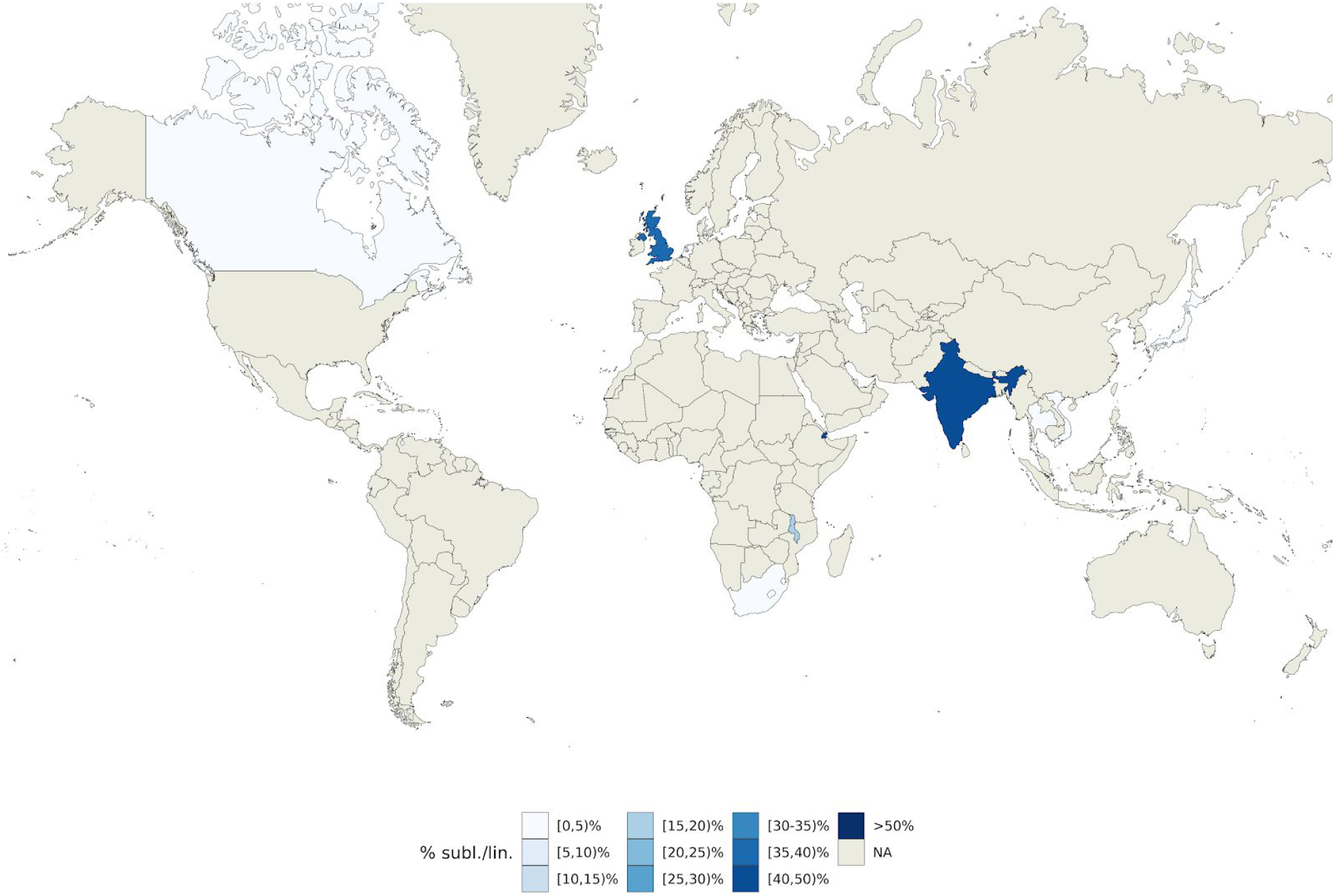
World map showing the geographic distribution of internal sub-lineage 1.1.2. Colors represent the percentage of 1.1.2 strains isolated in a given country with respect to all L1 strains isolated in such country.

To validate the geographic distribution of sub-lineages, we curated a dataset of 3,848 *Mtb* isolates sampled to reliably represent the entire population of TB patients in five countries (Azerbaijan, Bangladesh, Pakistan, South Africa and Ukraine, Suppl. File 5). In this dataset, we were able to identify 9 of the 30 new groups/sub-lineages described above. L1 and 1.1.2 isolates were found in three out of five countries (Azerbaijan, Pakistan and Ukraine) spanning two continents (Suppl. Fig. 14-15) supporting the hypothesis that L1.1.2 is a geographically unrestricted L1 sub-lineage. The candidate geographically restricted group 1.1.1.2.i1 from Malawi, was only additionally found in South Africa (Suppl. Fig. 15). L2 isolates were found in all five countries with 98% belonging to the modern-Beijing group (Suppl. Fig. 14 and 16). The modern-Beijing internal group 2.2.1.1.1.i3, corresponding to the Central Asia group ^9^, was the most prevalent L2 group in Ukraine and Azerbaijan and the second most prevalent group in Pakistan. This confirms 2.2.1.1.1.i3’s observed high Sdi (0.66) in the convenience training data and is in line with modern-Beijing’s transmissibility (next paragraph and ^16^). L3 isolates were found in four (Azerbaijan, Bangladesh, Pakistan, South Africa) of the five countries (Suppl. Fig. 14). Sub-lineage 3.1.1 was the most prevalent sub-lineage (Suppl. Fig. 17), but the two new L3 sub-lineages we describe (3.2.1 and 3.1.2) were also observed at low frequency (2.4-4.8%) in Bangladesh and Pakistan. L4 isolates and most commonly 4.1, 4.2.1.2.1.1 and 4.2.1.1.1.1.1.1 (that correspond to 4.1, 4.3/LAM and 4.10/PGG in the Stucki *et al*. ^10^ classification; Suppl. Fig. 18) were found in all five countries in line with their Sdi >0.67 and results on L4 transmissibility below.

### Differences in transmissibility between the Mtb global lineages

The observation that some lineages/sub-lineages are more geographically widespread than others raises the question of whether this results from differences in marginal transmissibility across human populations. On a topological level, we observed L2 and L3 phylogenies to be qualitatively different from those of L1 and L4 (Fig. 1-4): displaying a star-like pattern with shorter internal branches and longer branches near the termini. We confirmed this quantitatively by generating a single phylogenetic tree for all 9,584 L1-4 isolates and plotting cumulative branch lengths from root to tip for each main lineage (Suppl. Fig 19). Star-like topologies have been postulated to associate with rapid or effective viral or bacterial transmission *e*.*g*. a ‘super-spreading’ event in outbreak contexts ^19^. To compare transmissibility between the four lineages, we compared the distributions of terminal branch lengths expecting a skew towards shorter terminal branch lengths supporting the idea of higher transmissibility. We found L4 to have the shortest median terminal branch length, followed in order by L2, L3 and L1 (medians: 6.2×10^−5^, 8.2×10^−5^, 10.2 × 10^−5^, 17.5 × 10^−5^, respectively; all pairwise Wilcoxon rank sum tests significant P-value<0.001; Fig. 9). Shorter internal node-to-tip distance is a second phylogenetic correlate of transmissibility; the distribution of this measure across the four lineages revealed a similar hierarchy to the terminal branch length distribution (Suppl. Fig. 20). We also computed the cumulative distribution of isolates separated by increasing total pairwise SNS distance (Suppl. Fig. 21). The proportion of L4 isolates separated by <5 SNS was highest and followed by L2, L3 and L1, respectively. Thus, despite the topological similarities between the L3 and L2 trees, L3 was measured to be less transmissible by interrogating the topology of the trees.

**Fig. 9.**
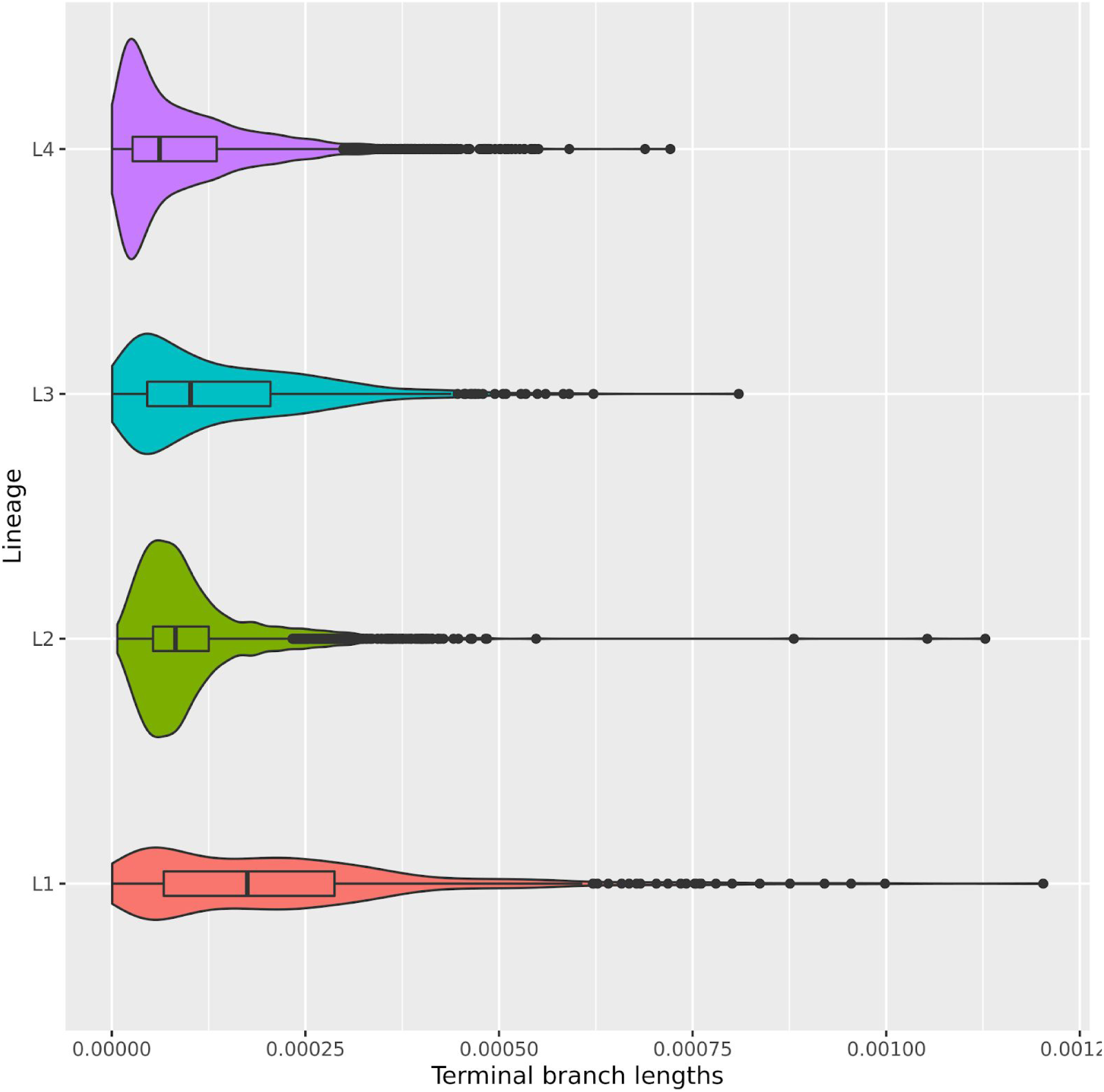
Violin plots showing the distributions of terminal branch lengths for the four global *Mtb* lineages (L1-L4). Wilcoxon Rank Sum tests were performed to test that two distributions were significantly different. Medians: 6.2×10^−5^ (L4), 8.2×10^−5^ (L2), 10.2 × 10^−5^ (L3), 17.5 × 10^−5^ (L1). Comparisons: L1 vs L2, L3 or L4 (p-value < 2.2×10^−16^); L2 vs L3 (p-value = 3.6 ×10^−6^), L2 vs L4 (p-value < 2.2×10^−16^); L3 vs L4 (p-value < 2.2×10^−16^).

## Discussion

Our results provide the most detailed picture of *Mtb*’s population structure to date and extend previous schemas ^8,*10*^ particularly for L1 and L3. We describe 7 and 4 new sub-lineages/internal groups, respectively, for these two lineages. The largest improvement was in increasing the resolution of sub-lineage 1.2 (5 subdivisions instead of 2), and in partitioning L3 into 4 groups since previously only a single group with one sub-lineage was characterized (3.1) for L3. For L2, we find limited internal diversity with the exception of major splits between the proto-, ancient and modern Beijing groups. Our schema supports 3 out of 4 groups previously described as either proto-Beijing or ancient-Beijing; the fourth unsupported group in our schema is Asia ancestral 3 previously proposed as a subdivision of ancient Beijing ^9^. We also identify a separate new sub-lineage of the ancient-Beijing group (2.2.1.2). Our results show no evidence for sub-lineages inside the modern-Beijing group, but support three internal groups (2.2.1.1.1.i1, 2.2.1.1.1.i2, 2.2.1.1.1.i3) one of which (Central Asia) was previously described by Shitikov *et al*.. Our analysis supports the existing schemas for L4 ^8,*10*^ and extends them by defining 11 new sub-lineages (and 5 internal groups). In particular, we improve the resolution of known sub-lineages, for instance 4.2.2, 4.2.1.1.1.2.1 and 4.2.1.1.1.2.2 (4.2, 4.6.1 and 4.6.2, respectively, in the Coll *et al*. classification) and define three new sub-lineages (4.2.1.2.1.2, 4.2.1.1.1.1.2 and 4.2.1.1.1.1.1.2) which increase the total number of L4 major sub-lineages from 7 (4.1-4.6 and 4.10 in the Stucki *et al*. classification) to 10.

We define a SNS barcode (95 SNS) that allows the rapid assignment of *Mtb* sub-lineage designations with detailed semantic and hierarchical sub-lineage naming schema from genomic data (.vcf files) ^8–10^. Along with this, we provide a software package and comparative tables / figures to facilitate the interchange between five SNS schemas and three *Mtb* sub-lineage naming systems ^8–10,20^. We expect these tools to facilitate the use of the expanded *Mtb* classification by scholars of *Mtb* evolution and public health practitioners alike for applications such as the rapid assessment of potential outbreaks and isolate triage for detailed phylogenetic analyses. Further, low level susceptibility to specific *Mtb* drugs has been reported for some lineages ^21,22^. Improving the resolution of classification at the sub-lineage level can allow for a better understanding of these findings and potentially a more refined prediction of drug resistance profiles.

We find evidence supporting the *Mtb*-human co-evolution hypothesis and its corollary of lineage differential adaptation^10^, including a spectrum of transmissibility across the four major *Mtb* lineages. We characterize L4 and L2 as the most transmissible and L1 as the least transmissible lineage using phylogenetic metrics. This is consistent with previous studies that have identified L2 sub-lineages as more transmissible than L1 in Vietnam ^16^ and Malawi ^17^. Additionally our data supports that at a global phylogenetic scale L4 is as or more transmissible than L2. This result is also in line with a larger number of geographically unrestricted ‘generalist’ L4 and L2 sub-lineages observed (compared with L3 or L1). We find two new candidate geographically restricted sub-lineages/internal groups, one of them from Malawi and one from South-East Asia. We also report evidence of a geographically unrestricted ‘generalist’ sub-lineage in L1 (1.1.2). L1 is the most ancestral of the four main *Mtb* lineages (L1-4), and given the evidence of low transmissibility on average across countries and sub-lineages, the finding of a potential generalist L1 sub-lineage requires further study and validation with additional data.

Our results raise the question of whether the geographically restricted sub-lineages represent cases of specialization or adaptation to specific human populations and whether the geographically unrestricted sub-lineages might have genetic features that allow them to spread efficiently in many different human populations. Although the phylogenies and validated geographic distribution are suggestive of differential adaptation, confirmation of this observation requires control for TB exposure and differences in contact networks using epidemiological data. A limitation of our analysis is the lack of this data and other patient and country level data, including human ancestry, recent migration and HIV prevalence. Nevertheless we find large differences in the geographic distribution of more than 20 *Mtb* sub-lineages across 51 countries and validated using systematically sampled data from these respective geographies. In the future, as additional patient metadata and *Mtb* sequence data becomes available, follow up analyses can confirm the geographic distribution of these sub-lineages. In summary, this work provides new insights into the population structure, biogeography and transmissibility of *Mtb* and demonstrates the use of this information to classify *Mtb* strains and investigate the relationships between population structure, pathogenicity and transmission.

## Methods

### Phenotypic and geographic location data

We compiled a dataset of antibiotic susceptibility and resistance data for 11,349 *Mtb* isolates using public databases (Patric, ReSeqTB) ^23,24^ and literature curation ^11,25–36^. A summary table of the data and the scripts used to generate it are available at https://github.com/farhat-lab/resdata. The isolate metadata including the country of isolation were downloaded using metatools_ncbi (https://github.com/farhat-lab/metatools_ncbi). Isolates were filtered according to the procedures described below, resulting in a dataset of 9,584 isolates. The full list of isolates and countries of isolation are available on Suppl. File 1. We used this dataset to determine the sub-lineages definitions and study the differences in transmissibility between the major *Mtb* lineages. In order to determine the geographic distribution of the *Mtb* sub-lineages we used a larger dataset of 17,432 *Mtb* isolates. The full list of isolates and countries of isolation are available on Suppl. File 4. Finally, in order to validate the geographic distribution of sub-lineages we used a dataset of 3,848 *Mtb* isolates from five countries and three continents (Azerbaijan, Bangladesh, Pakistan, South Africa and Ukraine, Suppl. File 5). These isolates come from epidemiological surveys designed to be representative of the entire population of TB patients of each one of these countries ^13^.

### Sequencing data analysis and lineage calling

Sequence read data for the isolates were downloaded from NCBI (the list of BioSamples is available on Suppl. File 1). We used an implementation of the pipeline proposed by Ezewudo *et al*. ^*37*^ to get the genetic variants that characterize any strain with respect to the *Mtb* reference strain H37Rv. For each isolate, identified by a BioSample, we downloaded all the associated Illumina sequencing runs and we ran PRINSEQ ^38^ to trim and filter the reads (using an average phred score threshold of 20). Then we ran Kraken ^39^ and discarded the isolates where less than 90% of the reads were assigned to *Mtb*. For this purpose we set up a custom Kraken database (to reduce the memory requirements of the default database) using sequences from the major *Mtb* lineages (NCBI Reference sequences: NC_009565.1, NC_000962.3, NC_017524.1, NC_002755.2, NC_021054.1).

Reads were then aligned to the H37Rv (NC_000962.3) reference genome using “BWA mem” ^40^. Duplicate reads were removed with Picard (http://broadinstitute.github.io/picard/). As an additional quality check the coverage of the *Mtb* genome was evaluated using samtools ^41^ and all isolates having a coverage of less than 95% of the *Mtb* genome with a depth of at least 10X were dropped. Genetic variants were called using Pilon ^42^. Lineage calls were made using the fast-lineage-caller (https://github.com/farhat-lab/fast-lineage-caller), using the Coll *et al*. ^8^ and Shitikov *et al*. ^9^ SNS schemes.

### Phylogenetic trees

In order to reduce the probability of having mixed isolates *(e*.*g*. isolates from different lineages/sub-lineages that infected the same patient), that could affect the determination of the tree of a given *Mtb* lineage, we computed the F2 lineage-mixture metric ^43^ and excluded the isolates that had scores greater than 0.5. We separately determined the phylogenetic trees for the pan-susceptible isolates (isolates that were susceptible to both isoniazid and rifampicin as well as all other antibiotics they were tested) and the resistant ones (isolates that were resistant to one or more antibiotics). To generate the trees we merged the .vcf files of the isolates of a given lineage with bcftools ^44^. Then we removed the repetitive, antibiotic resistance and low coverage regions (Suppl. File 6). We generated a multi-sequence .fasta alignment from the merged .vcf with vcf2phylip (version 1.5, https://doi.org/10.5281/zenodo.1257057) and we determined the phylogenetic tree with iqtree (version 1.6.10, automatic model selection, 1000 bootstraps).

### F_ST_ and PCA

The F_ST_ was calculated using the PopGenome R package (version 2.6.1) ^45^. PCA analysis was performed using the ade4 R package ^46^ (version 1.7-8).

### Sub-lineage definition

We defined the sub-lineages by evaluating: (1) the information of the tree topology, (2) the support (bootstrap), (3) the F_ST_ and (4) the results of a PCA analysis performed at each node. We considered a node as an informative one if: the support was greater than 95 (iqtree ultrafast bootstrap), the F_ST_ was greater than 0.33 and in the PCA analysis we were able to identify two groups (since our trees were binary trees). The threshold of the F_ST_ (0.33) we choose was already used to characterize sub-lineages in previous works ^10^ and the distributions of the F_ST_ calculated at each internal node of all lineages suggest it is a reasonable value to discriminate between *Mtb* sub-lineages (Suppl. Fig. 22). In a few situations (n = 3) we relaxed one of these constraints either because the node was the first split within a major lineage (split between L1.1 and L1.2, split between L4.1 and 4.2) or the group was already identified by previous studies and the results taken together were indicating that the group is valid (L4.2.1.2.1.1.i2). The scripts used to define the sub-lineages/internal groups are available at https://github.com/farhat-lab/mtb-popstruct-2020.

### Sub-lineage naming system

We assigned to all the sub-lineages their major lineage number (1, 2, 3 or 4). For each of the informative nodes in the tree (it is important to notice that we generated binary trees), we assigned a “.1” for the subtree having most of the descendants, while we assigned a “.2” for the subtree having the least descendants (exceptions to this rule were made for L4.1 and L2.1 to retain compatibility with the Coll *et al*. ^8^ SNS scheme). We stopped defining groups when a split was not sufficiently supported (ultrafast bootstrap < 95), the F_ST_ was less than 0.33 or the results of a PCA analysis were not conclusive (see also the Sub-lineage definition section). We also allowed for internal groups, *i*.*e*. if one or more ancestors of a given informative node were not informative we considered all the descendants of the informative node as members of an internal group of a given sub-lineage. We therefore added a “.i<n>” suffix to design such groups of strains as internal groups (*e*.*g*. 4.1.i1).

### Comparison of tree topologies

In order to compare the topologies of L1-4 trees we calculated the proportion of tree length as a function of the node level. We used a phylogenetic tree containing all L1-4 isolates. Starting from the MRCA node of each one of the L1-4 sets of isolates we iteratively looked for the children nodes (target nodes) and determined the average patristic distances between the target nodes and the MRCA node. Each one of the iterations represented a node level. We then normalized the resulting averaged distances by the maximum value of average patristic distance obtained. This algorithm was implemented in R using the ape ^47^ and phangorn ^48^ packages.

### A python lineage caller

We developed a python module (https://github.com/farhat-lab/fast-lineage-caller) that takes as input a .vcf file and returns the lineage / sub-lineage calls. We provide five SNS schemes for *Mtb*: Coll *et al*. ^8^ *(all lineages), the Shitikov et al*. ^9^ *(L2), Stucki et al*. ^*10*^ (L4, geographically bounded and unbounded sub-lineages), Lipworth *et al*. ^20^ *(Mtb* complex species, *Mtb* main lineages) and the one proposed here (L1-4).

### Simpson diversity index

The Simpson diversity index was calculated using the vegan (version 2.5.6) (https://cran.r-project.org/package=vegan) R package.

## Acknowledgements

We thank Karel Břinda for comments on the manuscript.

