## Supplementary Information for "Population structure, biogeography and transmissibility of *Mycobacterium tuberculosis*"

### Supplementary Figures

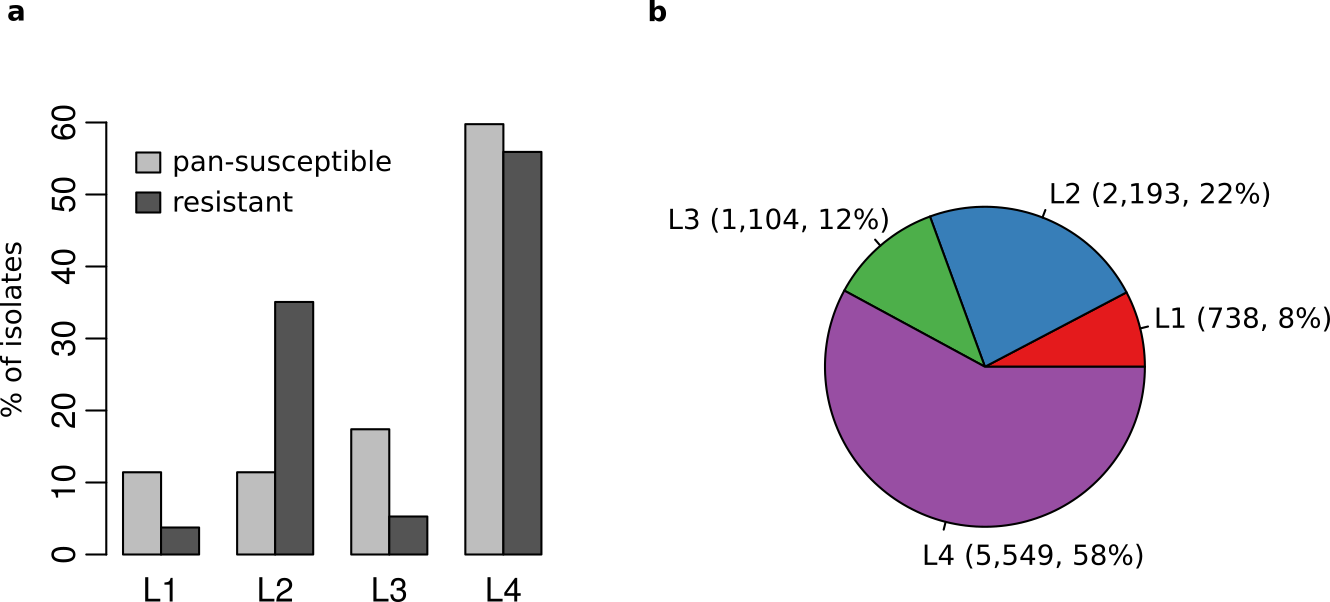

Suppl. Figure 1. **a** Bar chart showing the percentage of pan-susceptible or resistant isolates for each one of the four major *Mtb* lineages. **b** Pie chart showing the number of isolates present in our dataset for each one of the four major *Mtb* lineages (L1-4). Percentages relative to the total number of isolates of our dataset are also shown.

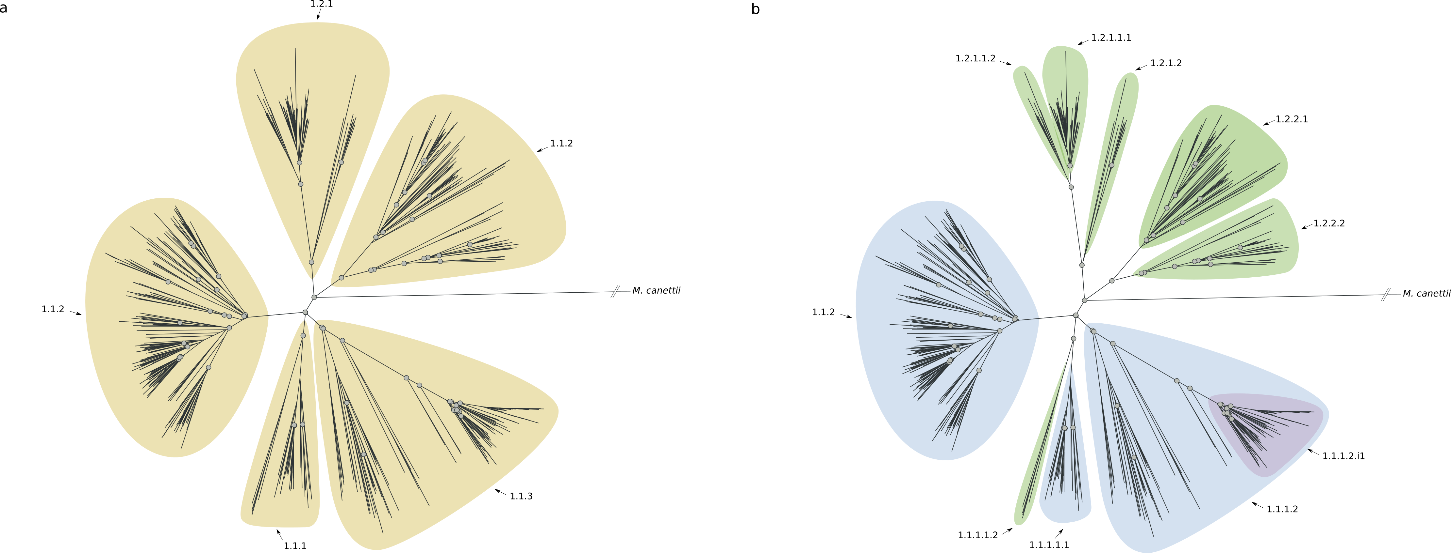

Suppl. Figure 2. **a** Phylogenetic tree reconstruction of L1. Yellow and orange areas define sub-lineages of L1 described by *Coll et al.* ^1^ **b** Phylogenetic tree reconstruction of L1. Colored areas define sub-lineages of L1 as described in this study (blue: sub-lineages that match those already described in the literature; green: sub-lineages described here; purple: internal sub-lineages).

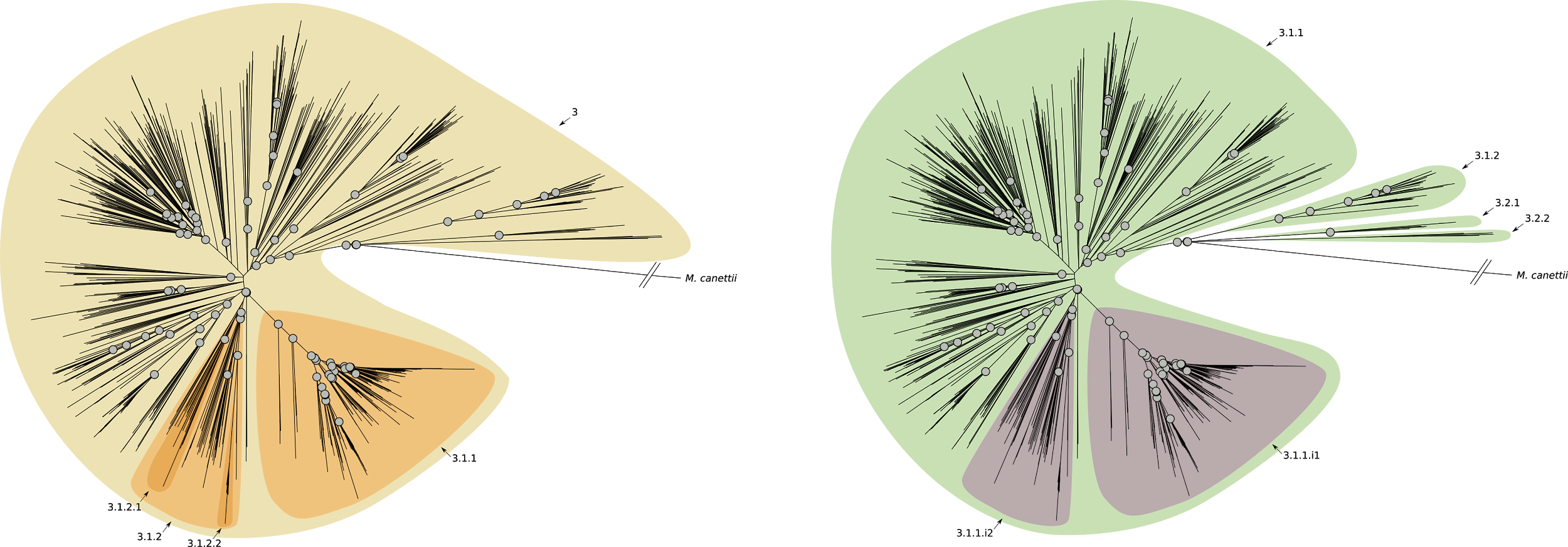

Suppl. Figure 3. **a** Phylogenetic tree reconstruction of L3. Yellow and orange areas define sub-lineages of L3 described by Coll *et al.* ^1^ ***b*** Phylogenetic tree reconstruction of L3. Colored areas define sub-lineages of L3 as described in this study (blue: sub-lineages that match those already described in the literature; green: sub-lineages described here; purple: internal sub-lineages).

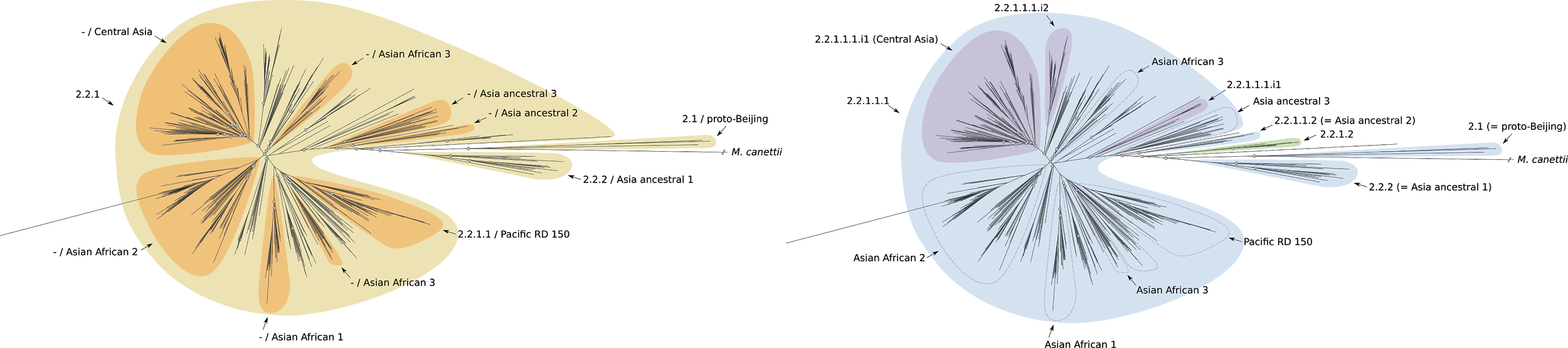

Suppl. Figure 4. **a** Phylogenetic tree reconstruction of L2. Yellow and orange areas define sub-lineages of L2 described by Coll *et al.* ^1^ or Shitikov *et al.* *^2^* ***b*** Phylogenetic tree reconstruction of L2. Colored areas define sub-lineages of L2 as described in this study (blue: sub-lineages that match those already described in the literature; green: sub-lineages described here; purple: internal sub-lineages).

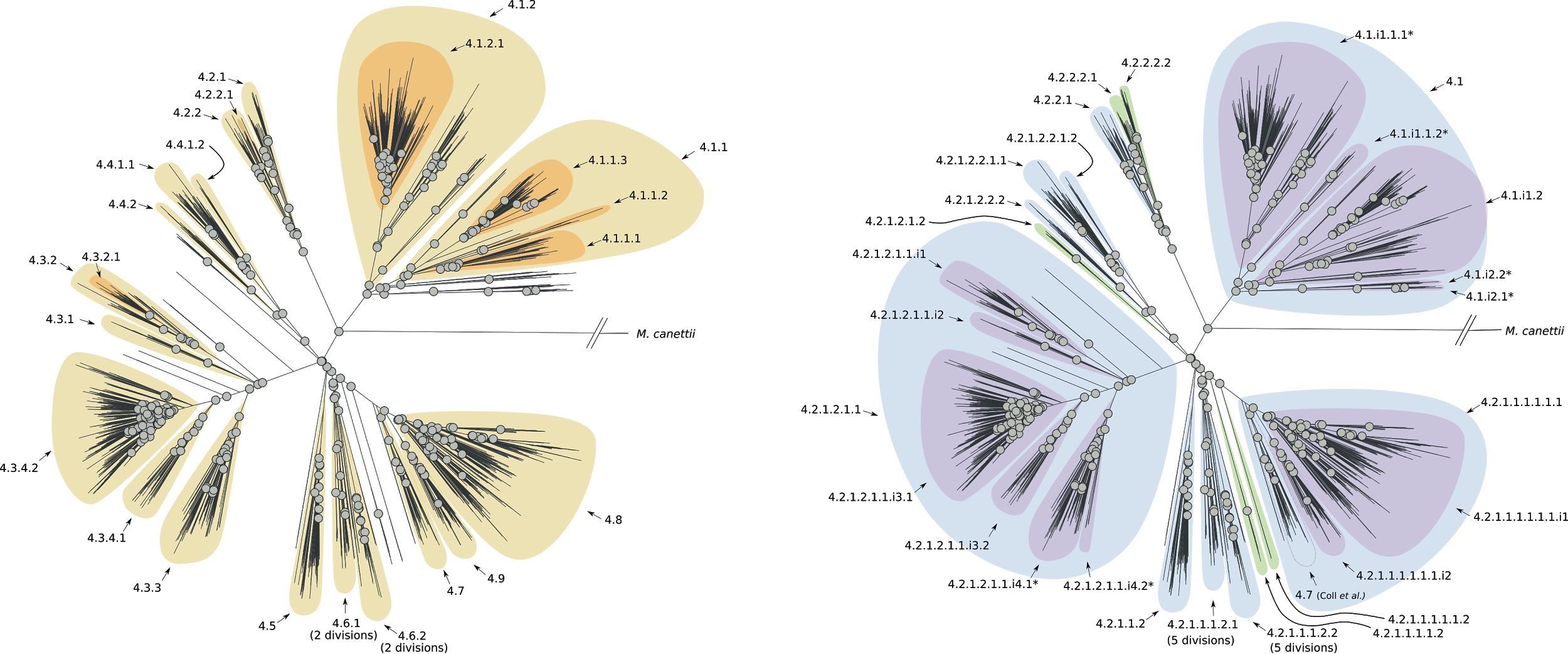

Suppl. Figure 5. **a** Phylogenetic tree reconstruction of L4. Yellow areas define sub-lineages of L4 described by Coll *et al.* ^1^ ***b*** Phylogenetic tree reconstruction of L4. Colored areas define sub-lineages of L4 as described in this study (blue: sub-lineages that match those already described in the literature; green: sub-lineages described here; purple: internal sub-lineages).

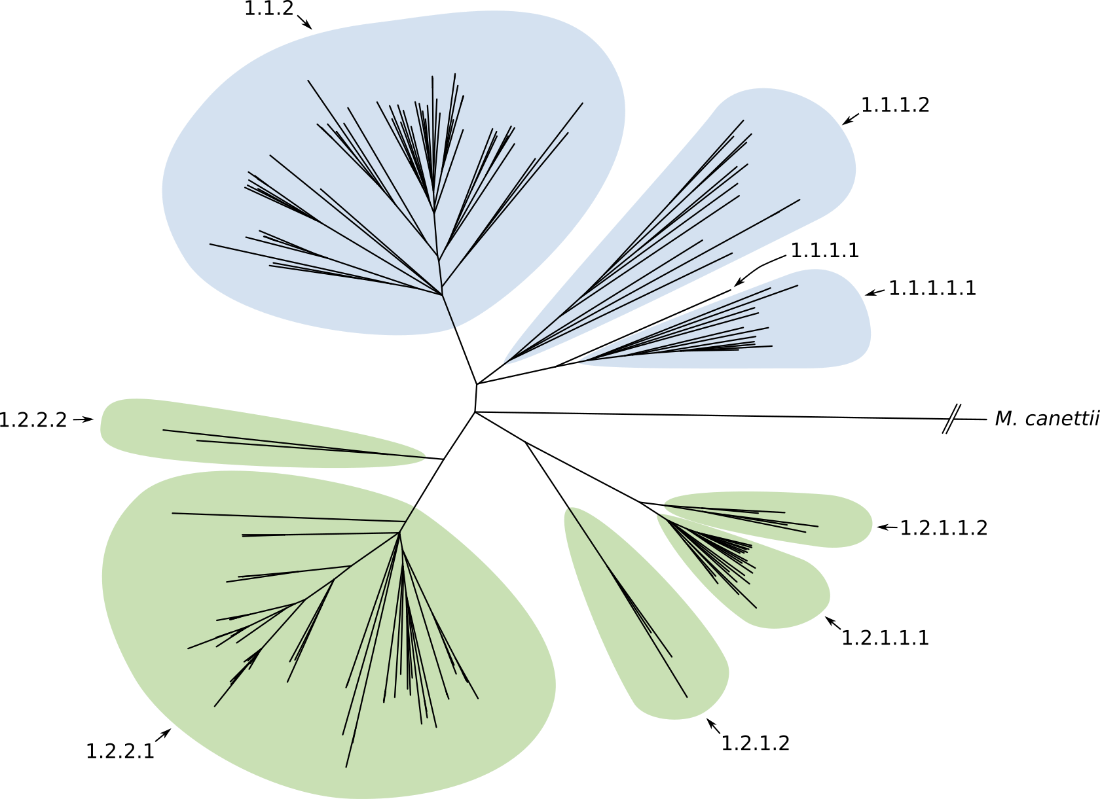

Suppl. Figure 6. Phylogenetic tree reconstruction of L1 resistant isolates (binary tree). Colored areas define sub-lineages of L1 as described in this study (blue: sub-lineages that match those already described in the literature; green: sub-lineages described here; purple: internal sub-lineages).

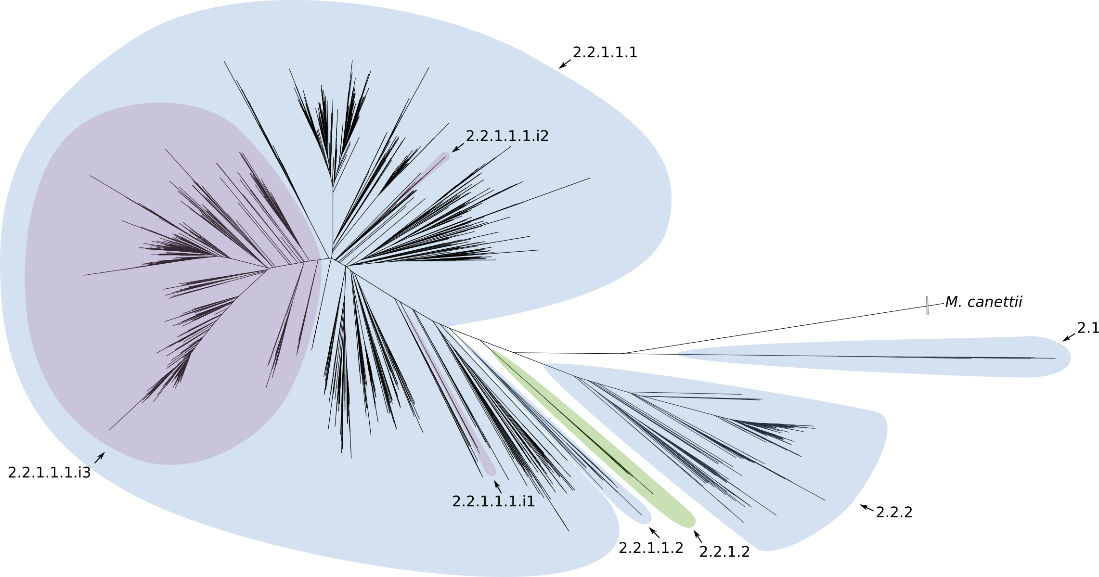

Suppl. Figure 7. Phylogenetic tree reconstruction of L2 resistant isolates (binary tree). Colored areas define sub-lineages of L2 as described in this study (blue: sub-lineages that match those already described in the literature; green: sub-lineages described here; purple: internal sub-lineages).

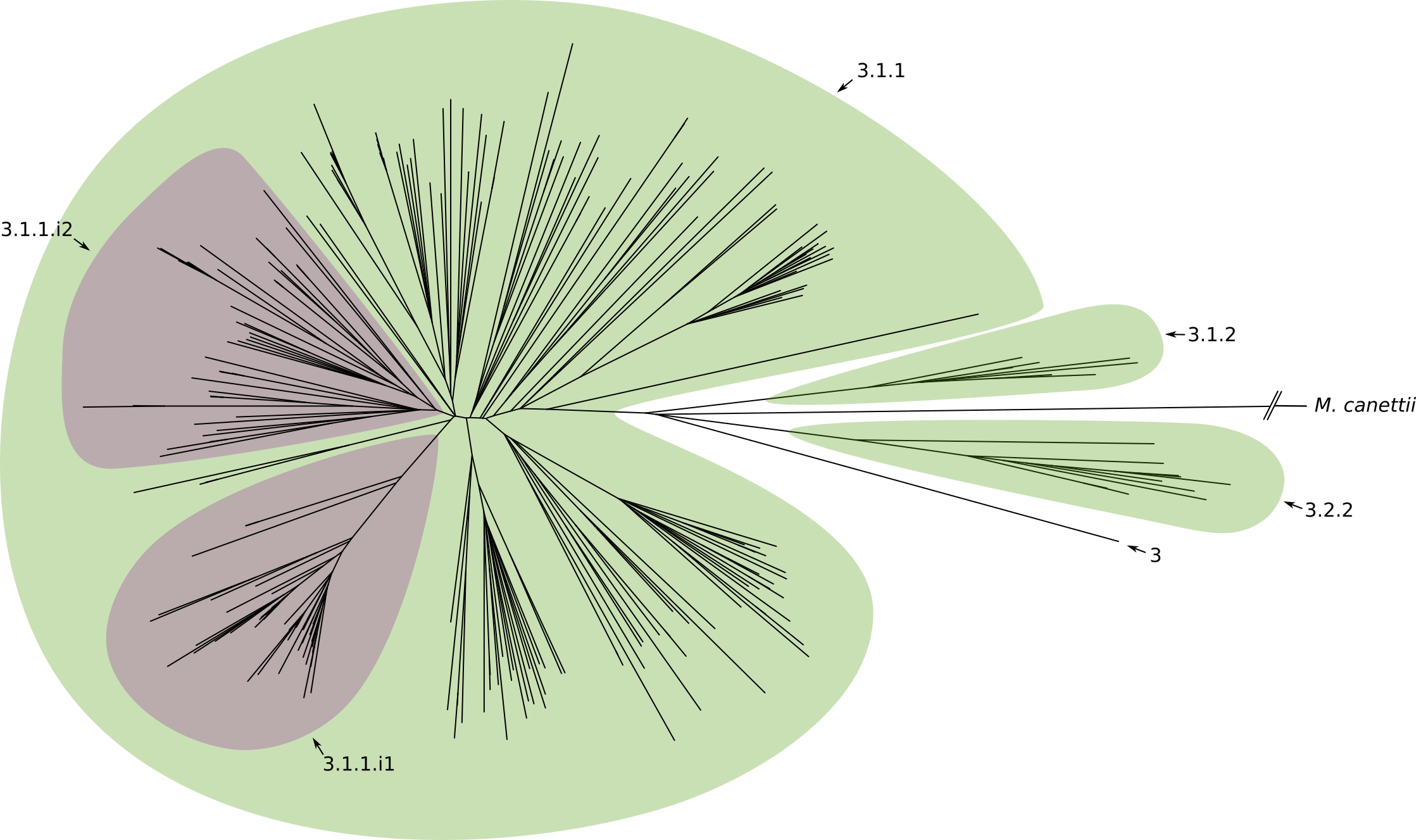

Suppl. Figure 8. Phylogenetic tree reconstruction of L3 resistant isolates (binary tree). Colored areas define sub-lineages of L3 as described in this study (blue: sub-lineages that match those already described in the literature; green: sub-lineages described here; purple: internal sub-lineages).

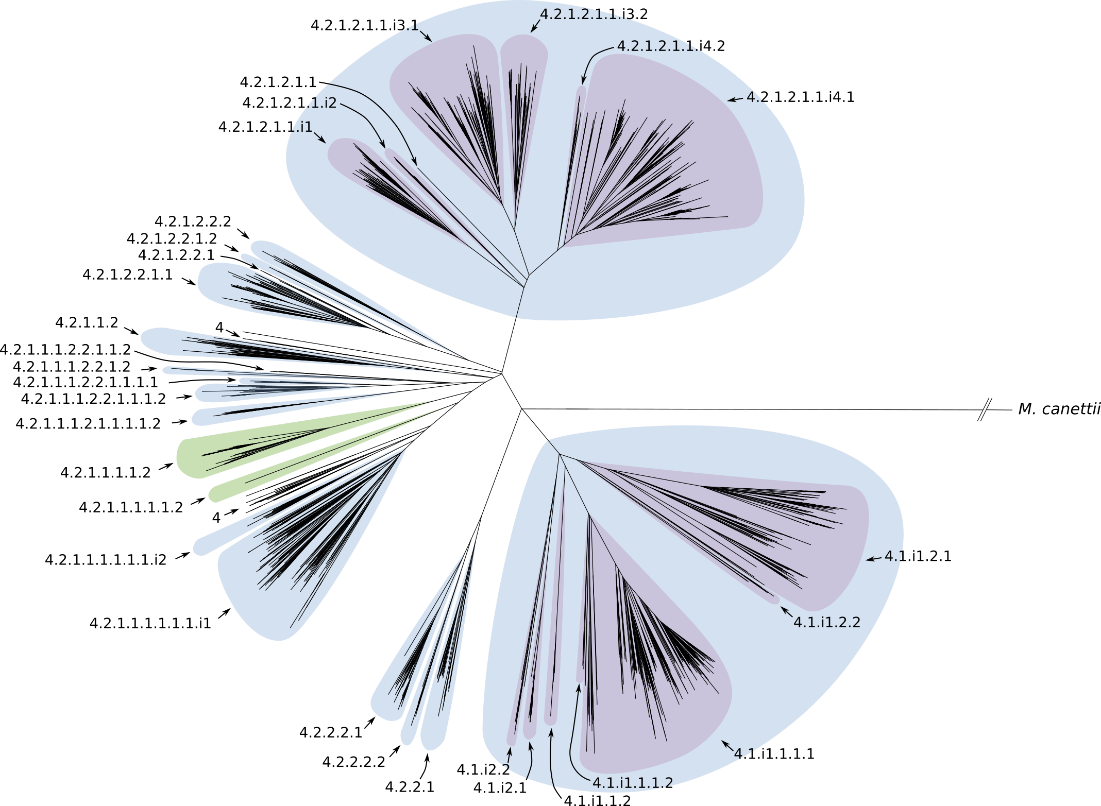

Suppl. Figure 9. Phylogenetic tree reconstruction of L4 resistant isolates (binary tree). Colored areas define sub-lineages of L4 as described in this study (blue: sub-lineages that match those already described in the literature; green: sub-lineages described here; purple: internal sub-lineages).

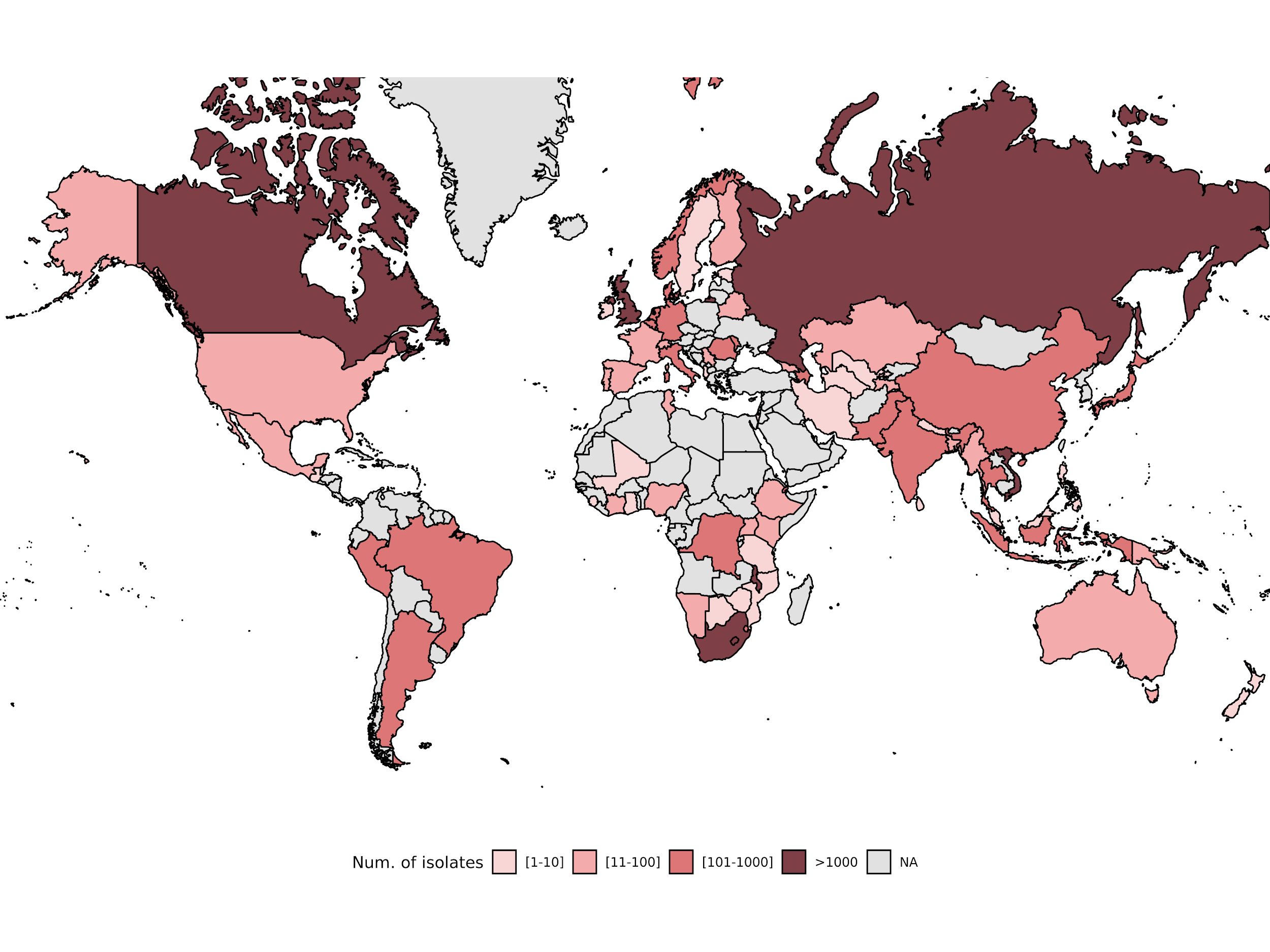

Suppl. Figure 10. Geographic distribution of the dataset of isolates used to study the biogeography of *Mtb* sub-lineages.

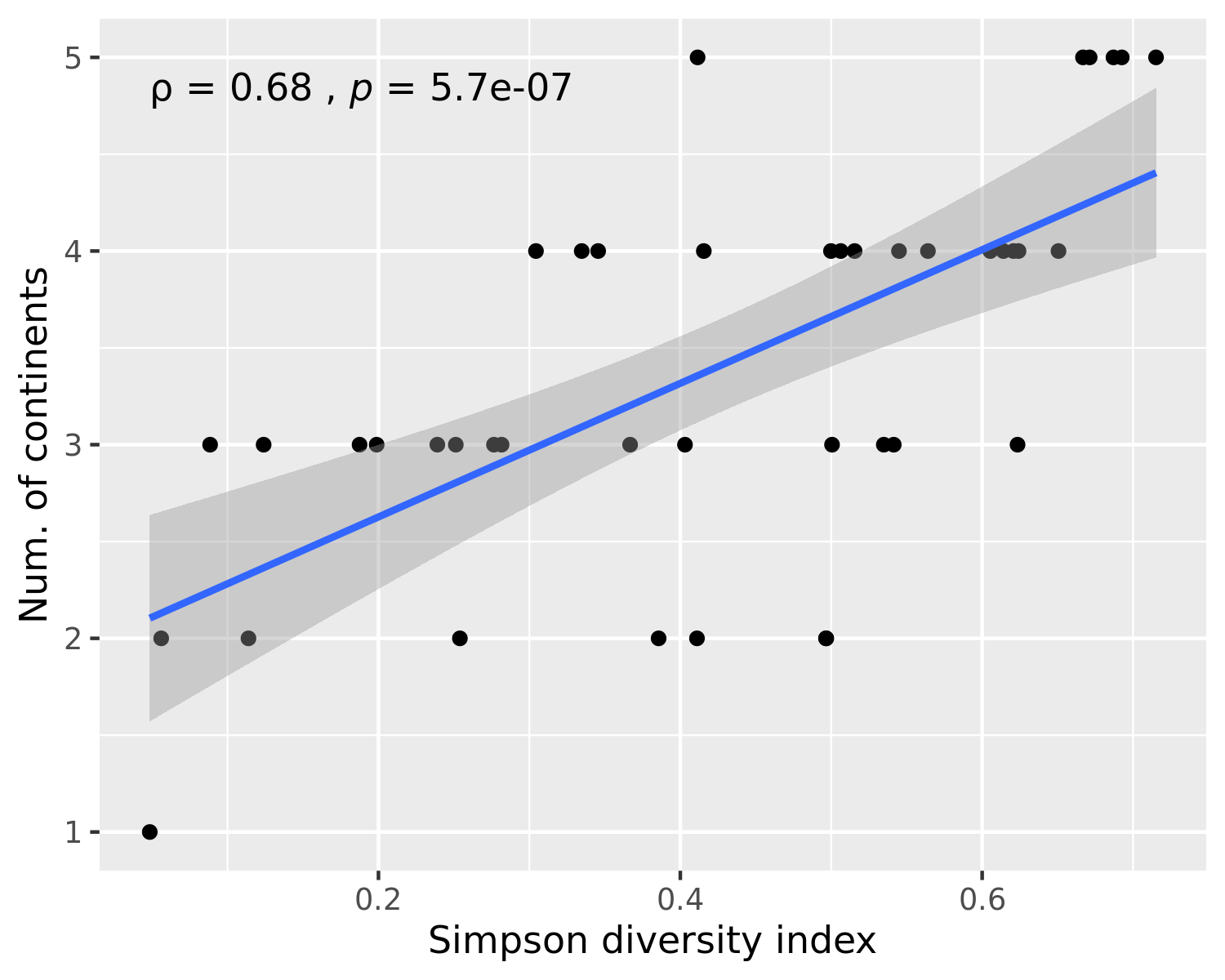

Suppl. Figure 11. Scatterplot that shows the relationship between the Simpson diversity index and the number of continents where a given sub-lineage has been found. The blue line shows a linear regression line and the grey area shows the 95% confidence interval. Spearman's rank correlation coefficient and the associated p-value are shown in the top-left part of the plot.

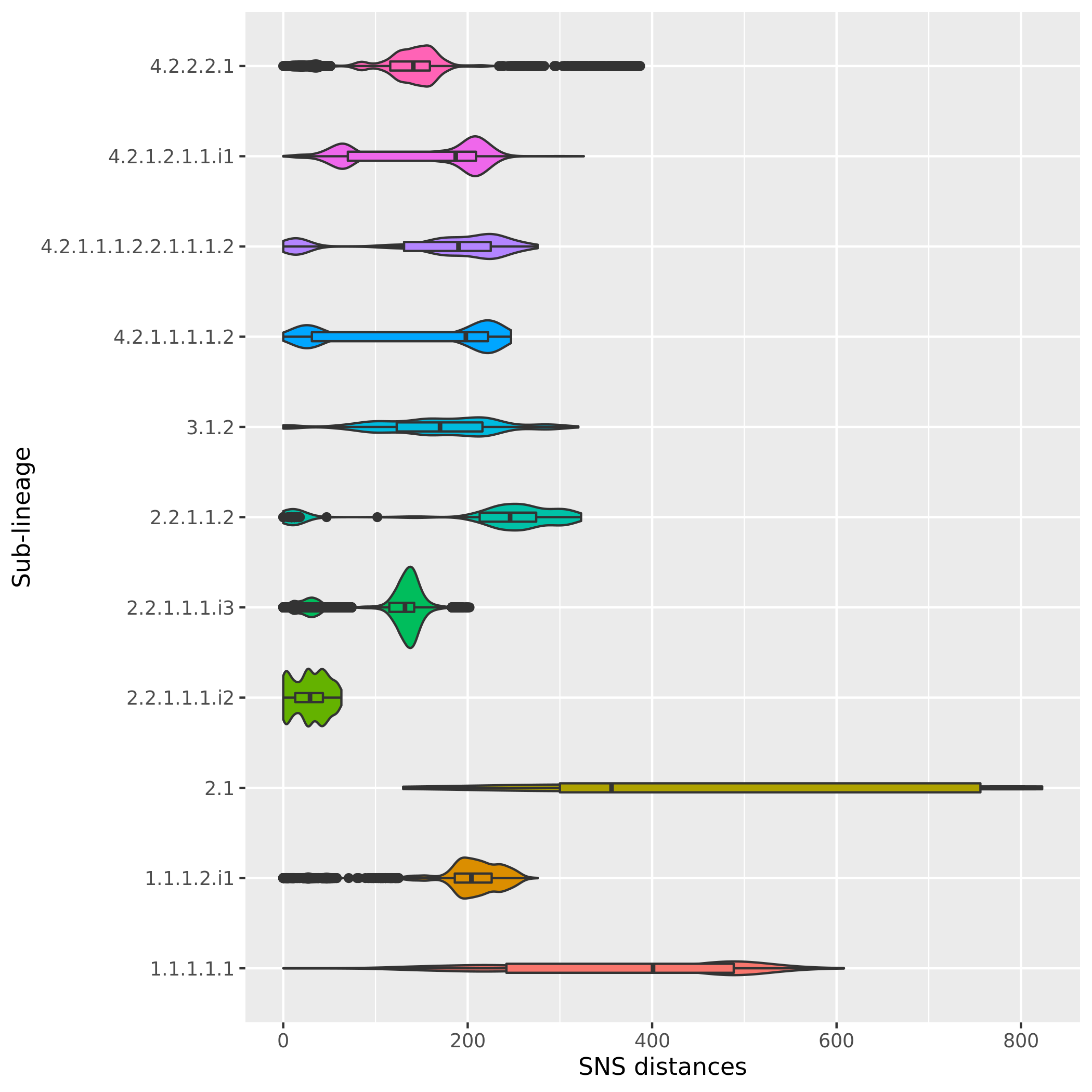

Suppl. Figure 12. Distributions of the pairwise SNS distances of the sub-lineages / internal groups that had Simpson diversity index < 0.28.

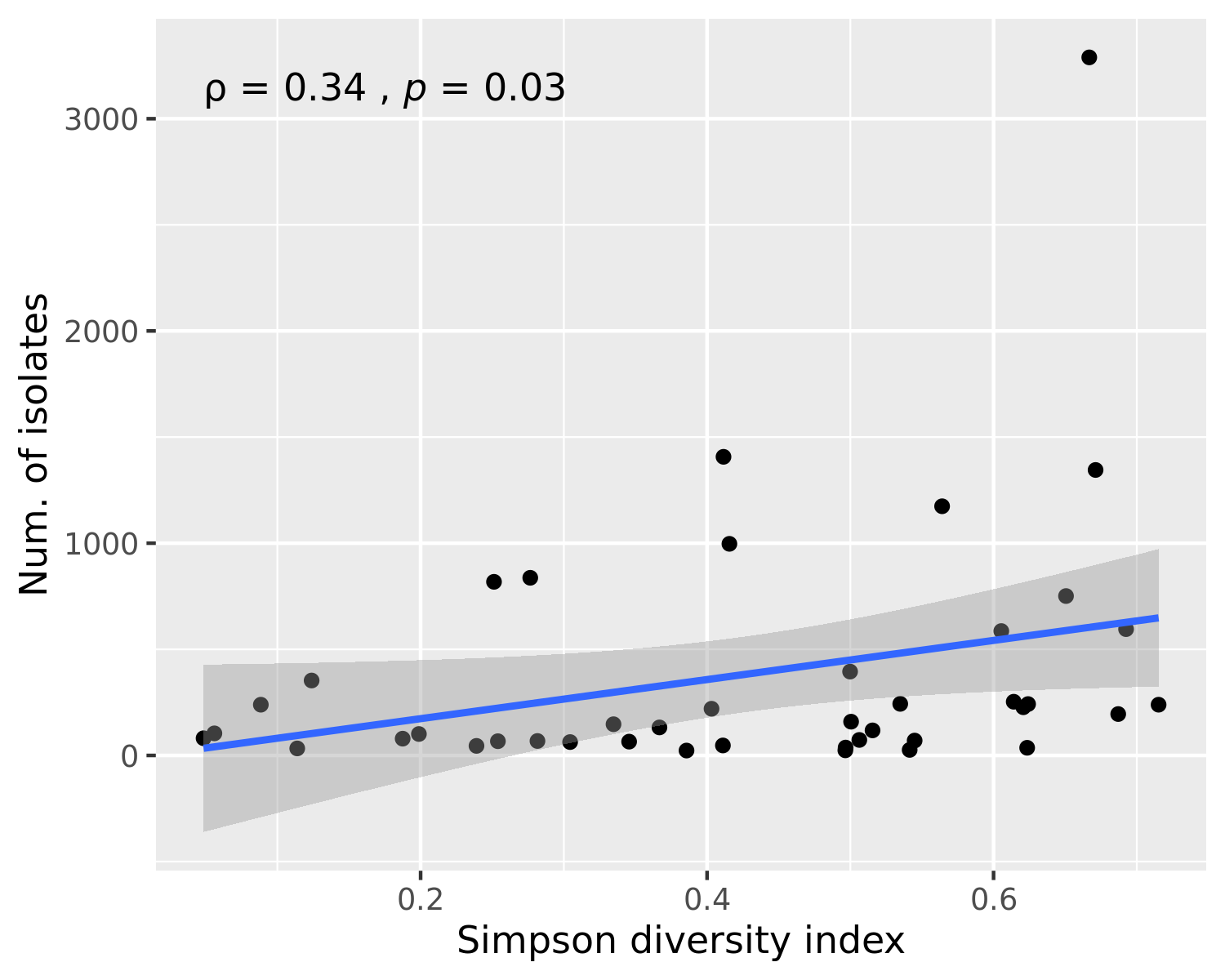

Suppl. Figure 13. Scatterplot that shows the relationship between the Simpson diversity index and the number of isolates of each sub-lineage. The blue line shows a linear regression line and the grey area shows the 95% confidence interval. Spearman's rank correlation coefficient and the associated p-value are shown in the top-left part of the plot.

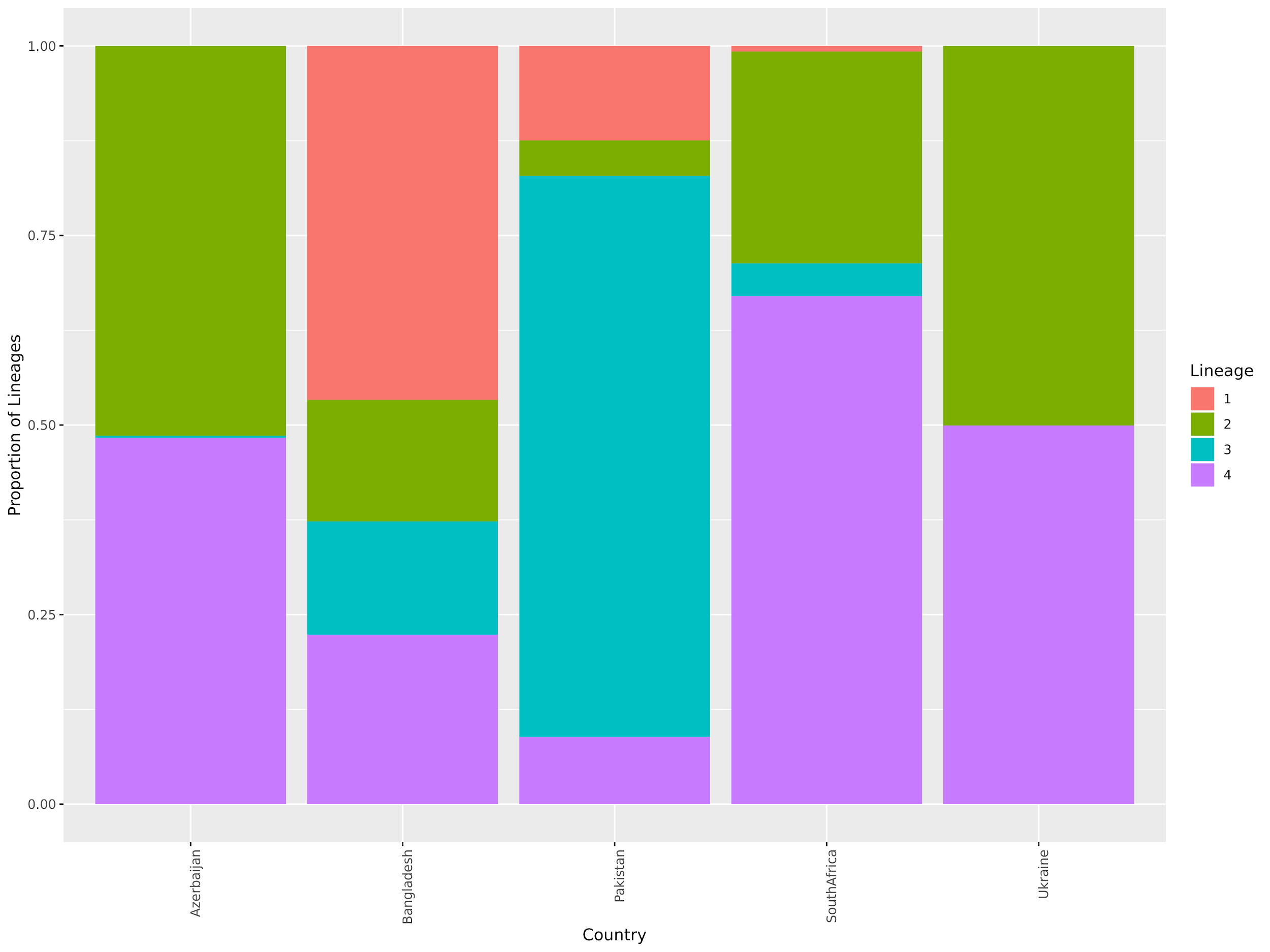

Suppl. Figure 14. Stacked bar chart showing the prevalence of different major lineages in five countries where isolates were randomly sampled.

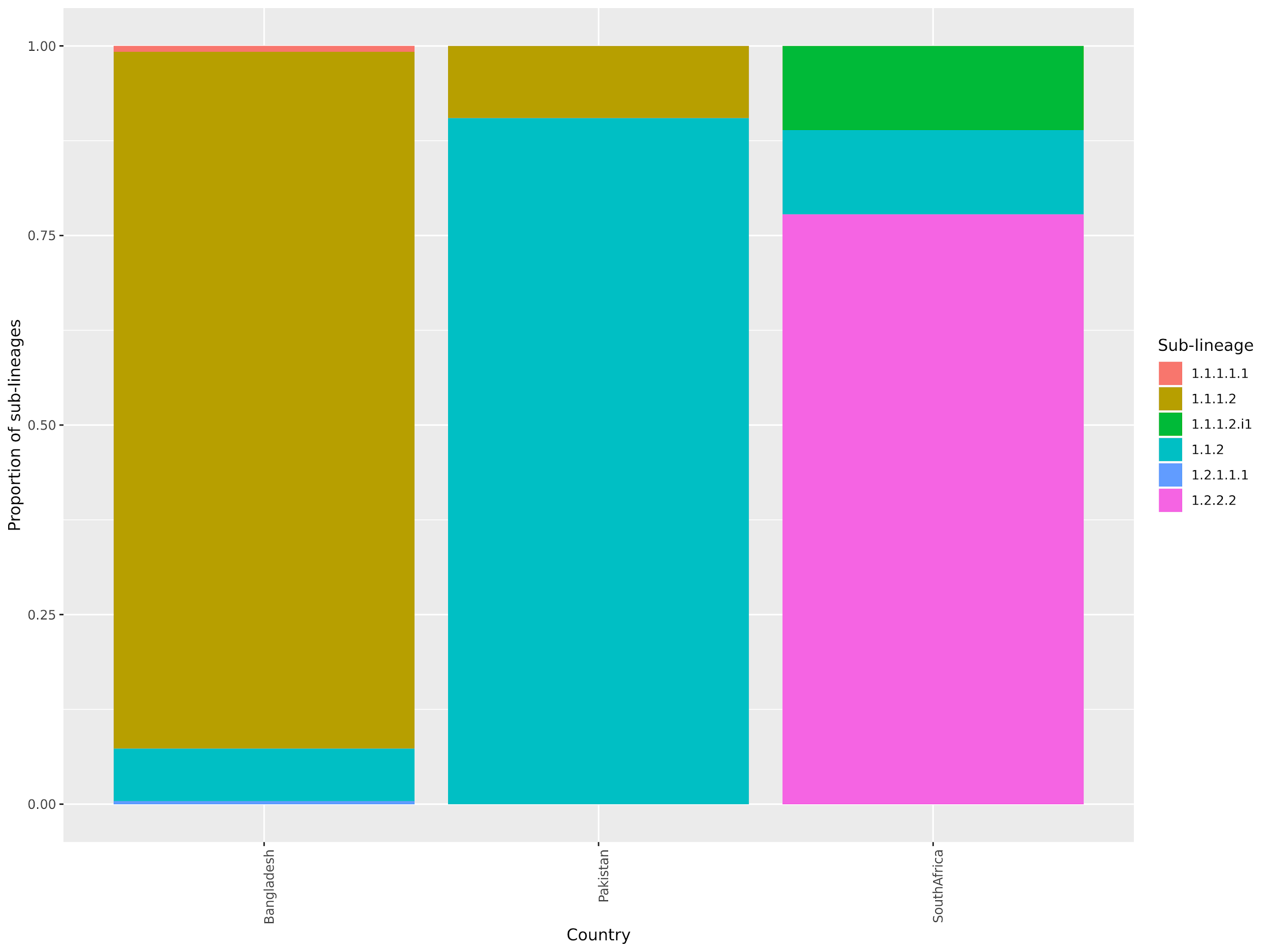

Suppl. Figure 15. Stacked bar chart showing the prevalence of different L1 sub-lineages in three countries where isolates were randomly sampled. No L1 isolates were found in Azerbaijan and Ukraine.

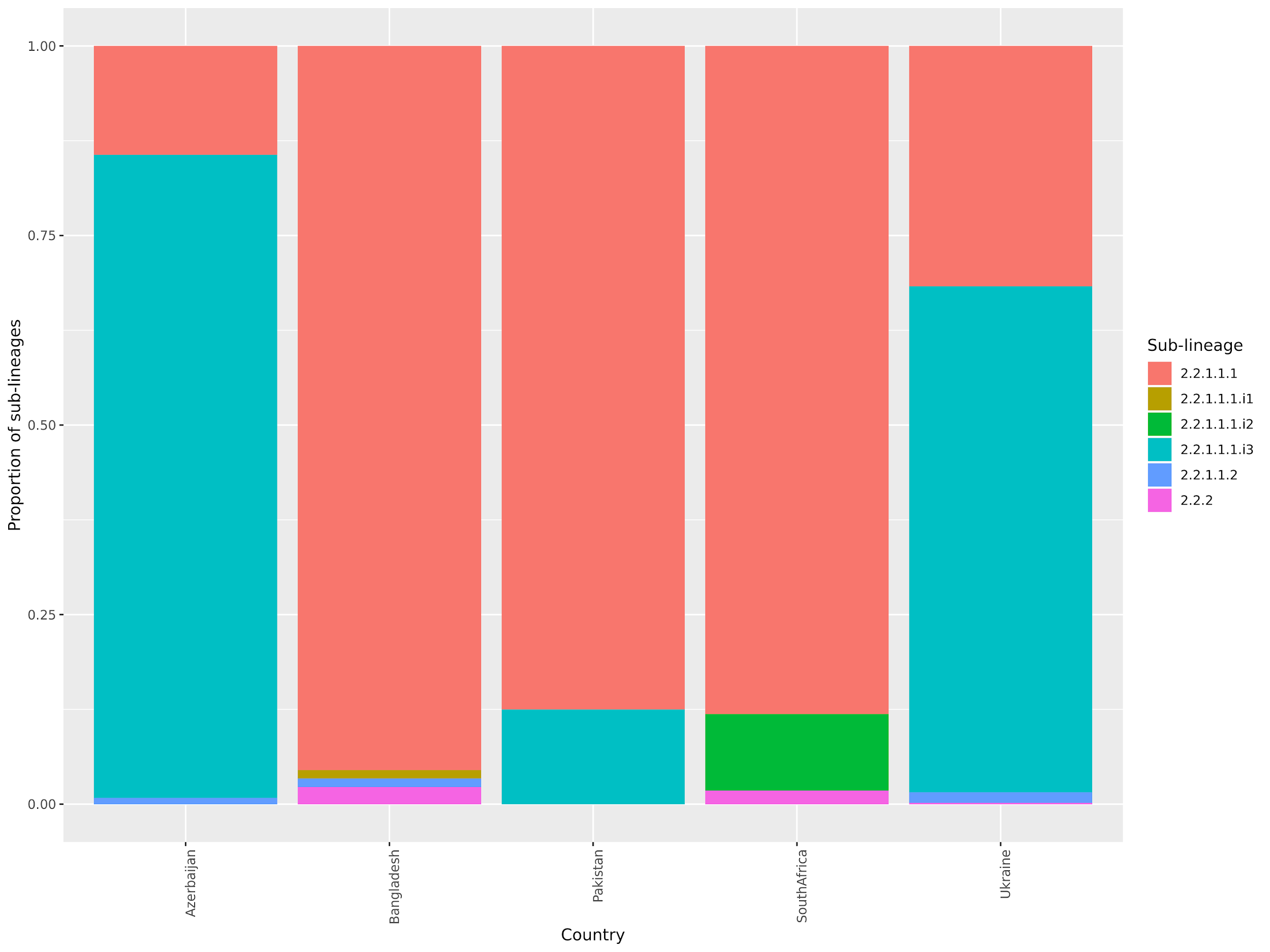

Suppl. Figure 16. Stacked bar chart showing the prevalence of different L2 sub-lineages in five countries where isolates were randomly sampled.

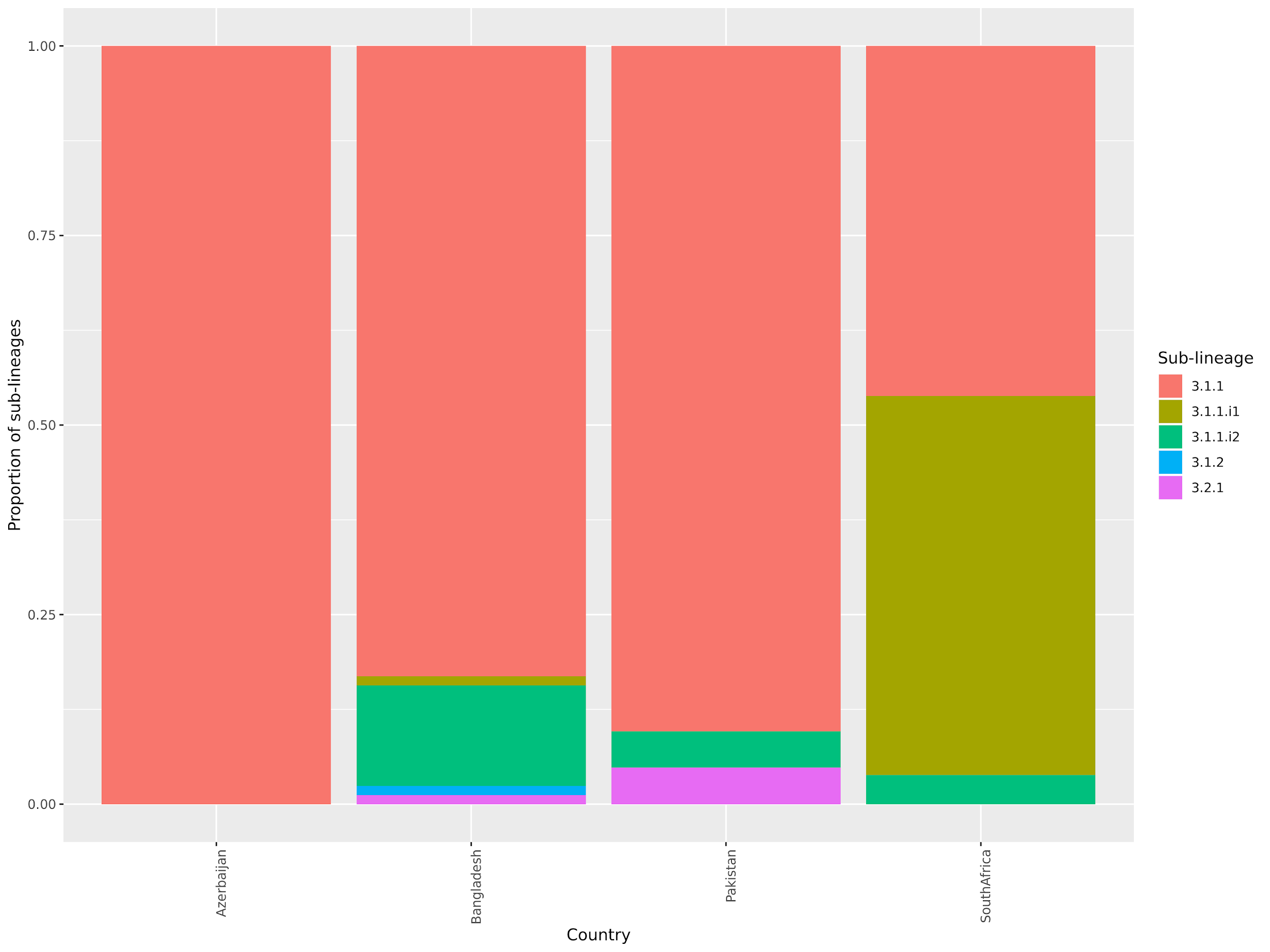

Suppl. Figure 17. Stacked bar chart showing the prevalence of different L3 sub-lineages in four countries where isolates were randomly sampled. No L3 isolates were found in Ukraine.

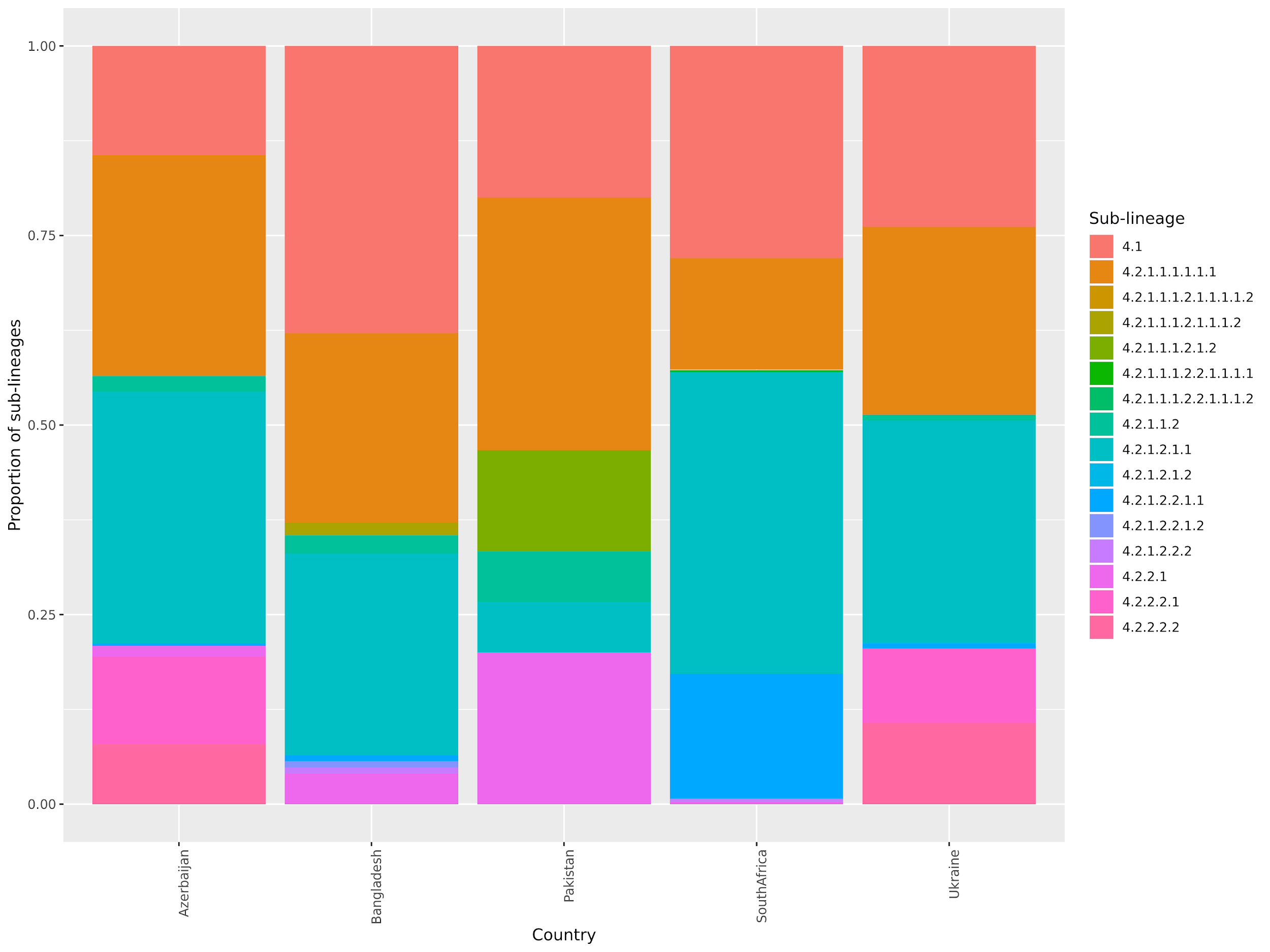

Suppl. Figure 18. Stacked bar chart showing the prevalence of different L4 sub-lineages in five countries where isolates were randomly sampled.

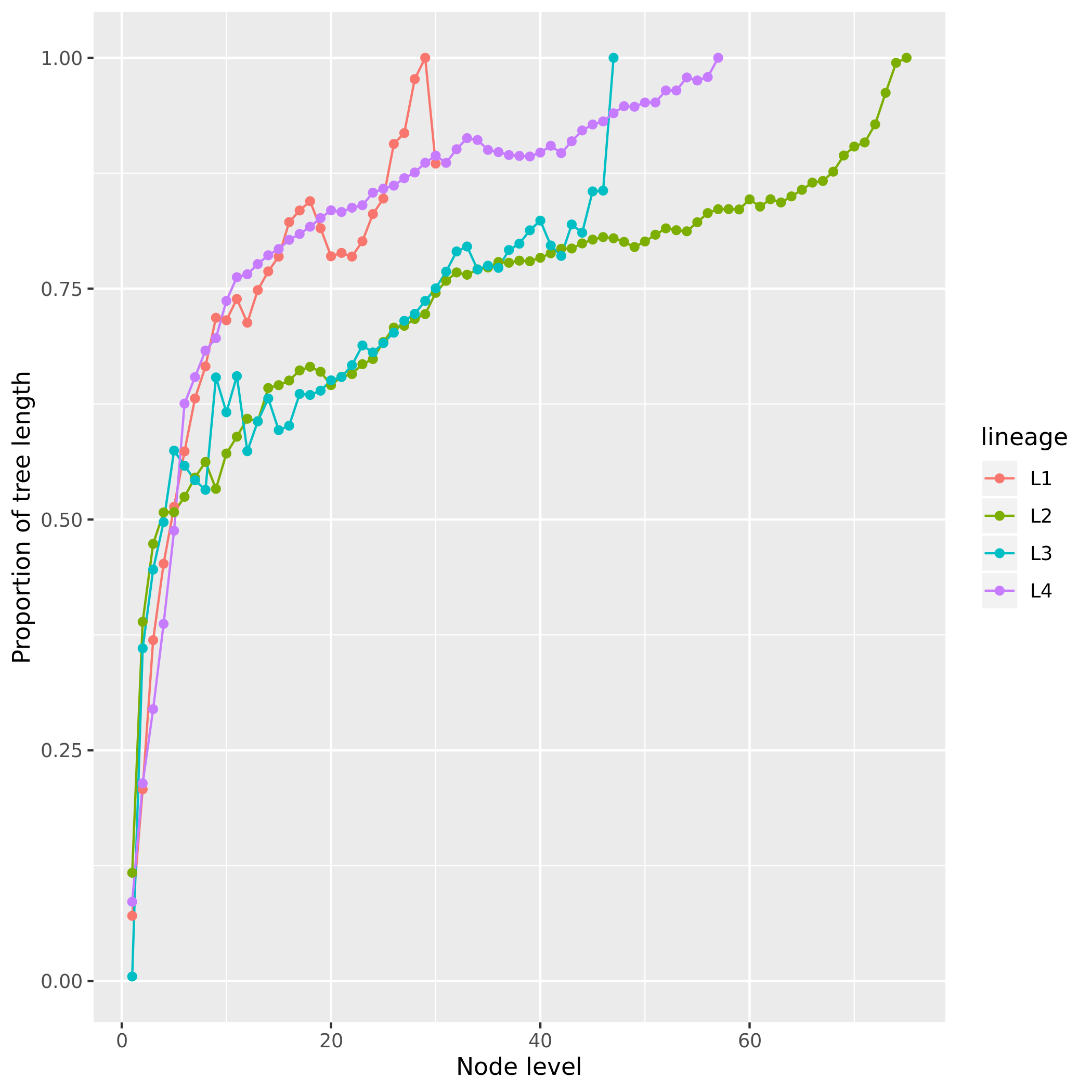

Suppl. Figure 19. Proportion of tree length as a function of node level. Colors identify different *Mtb* lineages.

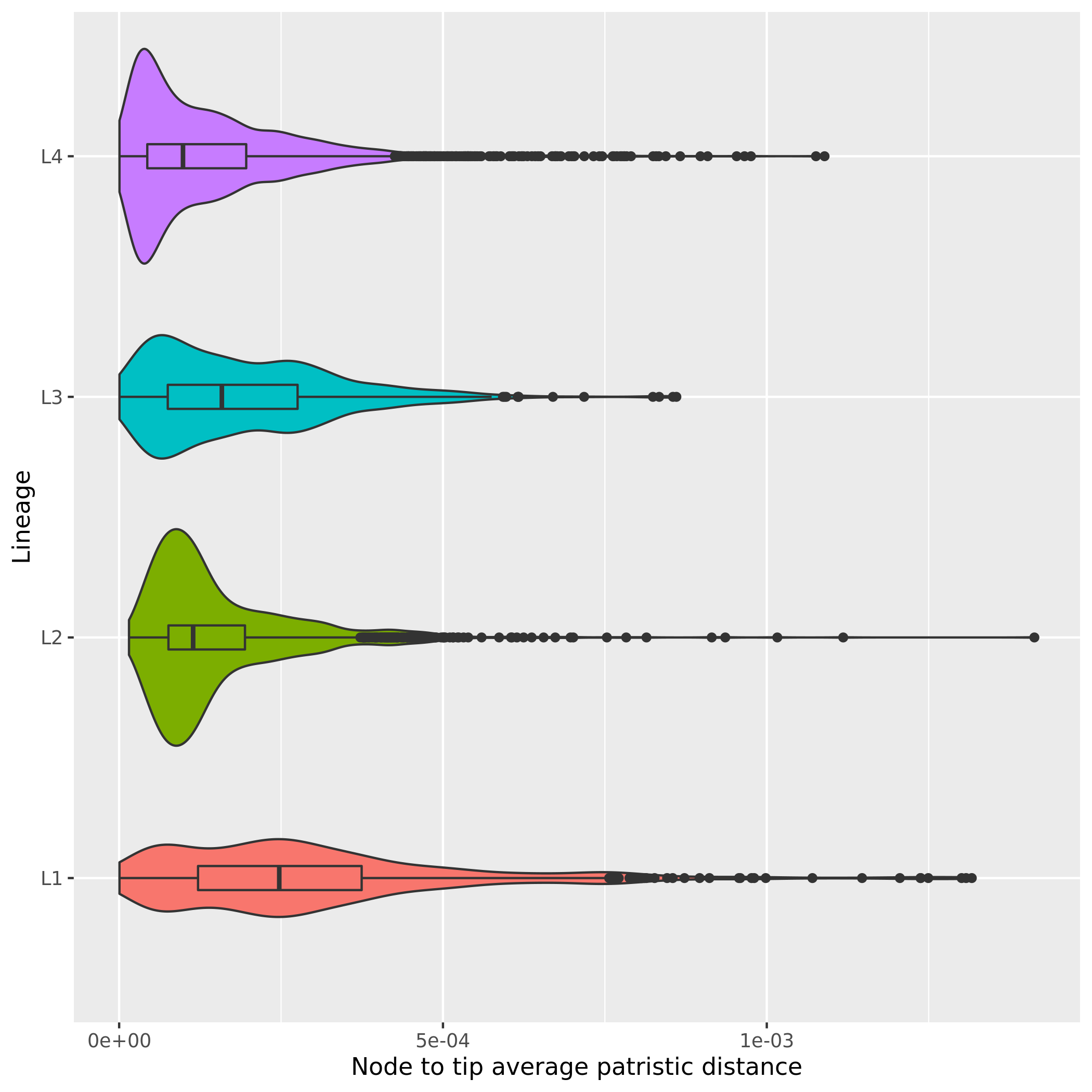

Suppl. Figure 20. Distribution of the node-to-tip distances for all internal nodes for all four *Mtb* lineages. Wilcoxon Rank Sum tests were performed to find if two distributions are significantly different. Medians: 9.85×10^-5^ (L4), 11.4×10^-5^ (L2), 15.8×10^-5^ (L3), 24.7×10^-5^ (L1). Comparisons: L1 vs L2 (p-value < 5.4×10^-60^); L1 vs L3 (p-value < 7.3×10^-23^); L1 vs L4 (p-value < 7.3×10^-99^); L2 vs L3 (p-value < 4.7×10^-9^); L2 vs L4 (p-value < 2.0×10^-24^); L3 vs L4 (p-value < 1.6×10^-37^).

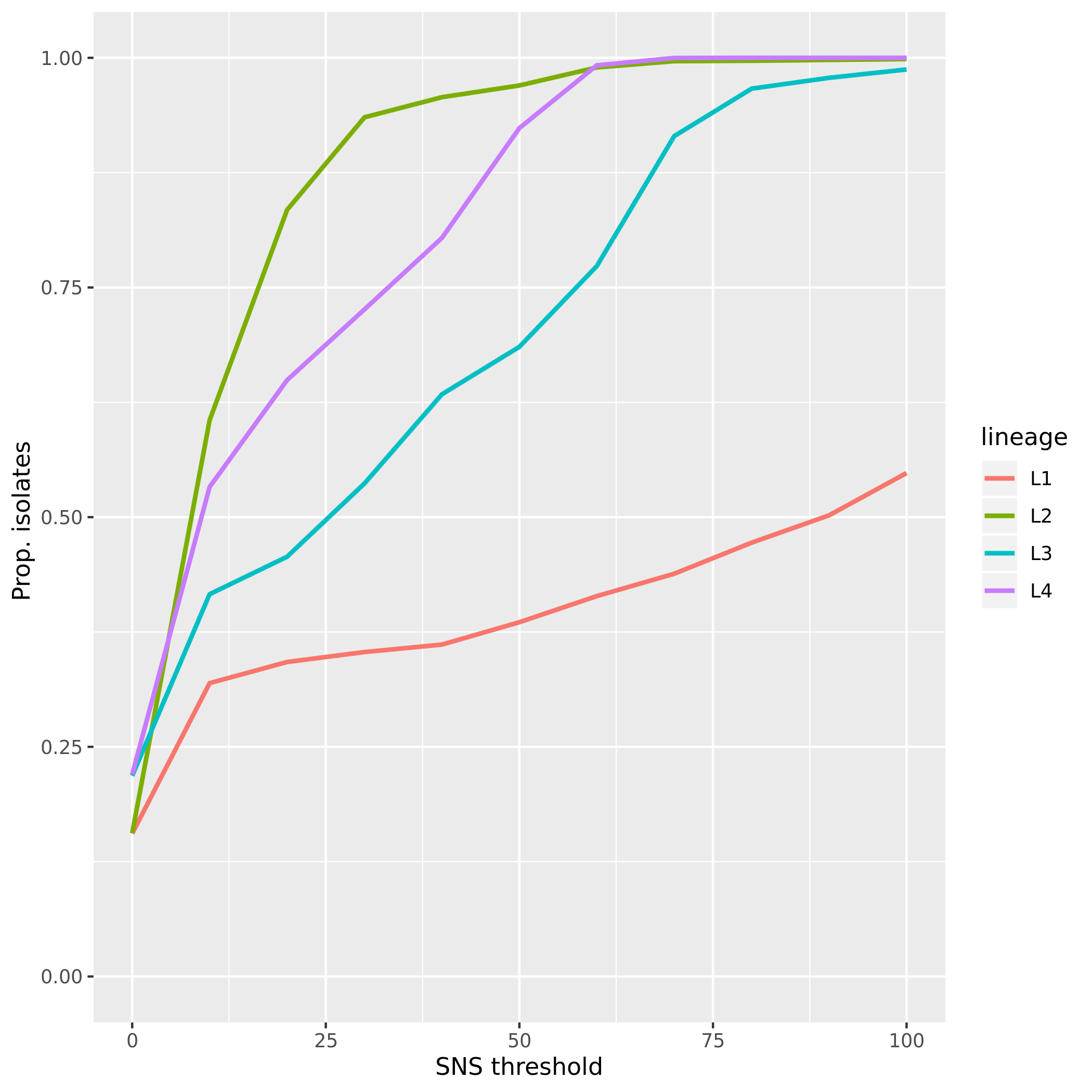

Suppl. Figure 21. Proportion of isolates from each one of the four major lineages (L1-4) that

belong to clusters defined at different pairwise SNS difference thresholds.

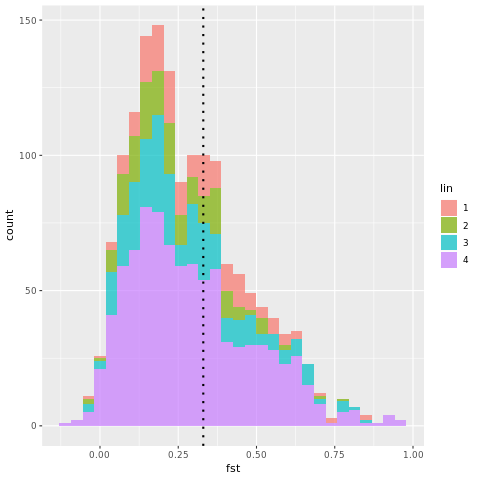

Suppl. Figure 22. Distribution of F_ST_ values calculated on all internal nodes of each of the phylogenetic trees of L1-4 (pan-susceptible isolates). The dotted line shows the threshold we chose to define two sub-lineages as distinct ones (0.33).

### Supplementary Tables

| **Sub-lineage** | **Simpson diversity index** |
| --- | --- |
| 4.6.2/Cameroon | 0.278 |
| 4.6.1/Uganda | 0.280 |
| 4.5 | 0.399 |
| 4.1.3/Ghana | 0.496 |

Suppl. Table 1. Simpson diversity index for the geographically restricted sub-lineages of L4 described by Stucki *et al.* ^3^

| **Sub-lineage** | **Simpson diversity index** |
| --- | --- |
| 4.10/PGG3 | 0.525 |
| 4.3/LAM | 0.553 |
| 4.1.2/Haarlem | 0.608 |

Suppl. Table 2. Simpson diversity index for the geographically unrestricted sub-lineages of L4 described by Stucki *et al.* ^3^

| **Sub-lineage** | **Simpson diversity index** | **Num. of isolates** | **Num. of countries** | **Num. of continents** | **Notes** |
| --- | --- | --- | --- | --- | --- |
| 2.2.1.1.1.i2 | 0.0484682 | 81 | 4 | 1 | Outbreak? |
| 1.1.1.2.i1 | 0.0560281 | 104 | 2 | 2 |  |
| 4.2.2.2.1 | 0.0884438 | 239 | 14 | 3 |  |
| 4.2.1.1.1.1.2 | 0.1138659 | 33 | 3 | 2 |  |
| 1.1.1.1.1 | 0.1239076 | 353 | 7 | 3 |  |
| 2.1 | 0.1874700 | 79 | 10 | 3 |  |
| 2.2.1.1.2 | 0.1988040 | 101 | 12 | 3 |  |
| 3.1.2 | 0.2390123 | 45 | 5 | 3 |  |
| 4.2.1.2.1.1.i1 | 0.2512359 | 818 | 18 | 3 |  |
| 4.2.1.1.1.2.2.1.1.1.2 | 0.2539541 | 67 | 7 | 2 | 4.6.1/Uganda |
| 2.2.1.1.1.i3 | 0.2765438 | 837 | 15 | 3 | Central Asia |

Suppl. Table 3. Summary table of the geographic distribution of sub-lineages / internal groups having Simpson diversity index < 0.28 (which corresponds to the lowest Simpson diversity index value for known geographically restricted sub-lineages)

| **Sub-lineage** | **Simpson diversity index** | **Num. of isolates** | **Num. of countries** | **Num. of continents** | **Notes** |
| --- | --- | --- | --- | --- | --- |
| 4.2.1.1.1.1.1.1.i2 | 0.7152186 | 239 | 21 | 5 | Corresp. to 4.9 / Internal group of 4.10/PGG3 |
| 4.2.1.2.1.1.i4.1 | 0.6925104 | 595 | 31 | 5 | Internal group of 4.3/LAM |
| 4.2.1.2.1.1.i3.2 | 0.6870217 | 195 | 19 | 5 | Internal group of 4.3/LAM |
| 4.1.i1.1.1.1 | 0.6712246 | 1345 | 33 | 5 | Internal group of 4.1/ corresp to 4.1.2/Haarlem |
| 2.2.1.1.1 | 0.6668162 | 3289 | 24 | 5 | ~ modern Beijing |
| 4.1.i1.2.1 | 0.6505840 | 751 | 24 | 4 |  |
| 2.2.2 | 0.6241035 | 242 | 14 | 4 |  |
| 1.2.1.1.2 | 0.6234568 | 36 | 6 | 3 |  |
| 4.2.2.1 | 0.6207767 | 227 | 20 | 4 |  |
| 1.1.2 | 0.6140699 | 253 | 7 | 4 |  |
| 4.2.1.2.2.1.1 | 0.6053944 | 586 | 23 | 4 | Corresp. to 4.4.1.1 |

Suppl. Table 4. Summary table of the geographic distribution of sub-lineages / internal groups having Simpson diversity index > 0.6 (which corresponds to the highest Simpson diversity index value for known geographically unrestricted sub-lineages)
